# Interspecies surfactants serve as public goods enabling surface motility in *Pseudomonas aeruginosa*

**DOI:** 10.1101/2024.01.03.573969

**Authors:** Delayna L. Warrell, Tiffany M. Zarrella, Christopher Machalek, Anupama Khare

**Affiliations:** Laboratory of Molecular Biology, Center for Cancer Research, National Cancer Institute, National Institutes of Health, Bethesda, MD, USA; Postdoctoral Research Associate Training Program, National Institute of General Medical Sciences, National Institutes of Health, Bethesda, MD, USA; Current address: Department of Biology, Georgetown University, Washington, DC, USA

**Keywords:** *Pseudomonas aeruginosa*, *Staphylococcus aureus*, surfactants, motility, polymicrobial interactions, phenol-soluble modulins

## Abstract

In most natural environments, bacteria live in polymicrobial communities where secreted molecules from neighboring species alter bacterial behaviors including motility, but such interactions are understudied. *Pseudomonas aeruginosa* is a motile opportunistic pathogen that exists in diverse multispecies environments such as the soil and is frequently found in human wound and respiratory tract co-infections with other bacteria including *Staphylococcus aureus*. Here we show that *P. aeruginosa* can co-opt secreted surfactants from other species for flagellar-based surface motility. We found that exogenous surfactants from *S. aureus*, other bacteria, and interkingdom species enabled *P. aeruginosa* to switch from swarming to an alternative surface spreading motility on semi-solid surfaces and allowed for the emergence of surface motility on hard agar where *P. aeruginosa* was otherwise unable to move. This motility was distinct from the response of other motile bacteria in the presence of exogenous surfactants. Mutant analysis indicated that this *P. aeruginosa* motility was similar to a previously described mucin-based motility, ‘surfing’, albeit with divergent regulation. Thus, our study demonstrates that secreted surfactants from the host as well as neighboring bacterial and interkingdom species act as public goods facilitating *P. aeruginosa* flagella-mediated surfing-like surface motility, thereby allowing it to access different environmental niches.

## INTRODUCTION

Bacteria are often found in dynamic, complex communities where interspecies secreted factors influence bacterial behaviors, such as antibiotic resistance (1–3), biofilm formation (4–7), and motility (8–10). However, many of the polymicrobial interactions leading to the modification of these pathogenic traits have not been elucidated. The study of pairwise interactions has the potential to reveal behavioral changes that occur within communities as well as the underlying molecular mechanisms (11, 12).

Motility allows bacterial cells to disperse and migrate to new niches for nutrients, seek out prey, and adapt to new environments, and modulation of motility behaviors due to ecological interactions is thus likely to be advantageous for bacterial fitness. Several examples of changes in bacterial motility in mixed cultures compared to monocultures have been previously reported, including the inhibition of motility via secreted volatiles (9), and emergence of motility in co-culture conditions where the monocultures did not move. Such emergent motility includes social spreading between two non-motile species (13), exploration by non-motile bacteria upon sensing fungal volatiles (8), and non-motile bacteria inducing motility in a neighboring species, and then hitchhiking on the moving population (14).

*Pseudomonas aeruginosa* is a social, motile microbe that typically inhabits polymicrobial environments in soil, wounds, and persistent infections of the respiratory tracts of people with cystic fibrosis (CF) (15–17). Motility in *P. aeruginosa* is intricately linked with virulence regulation, antibiotic resistance, and biofilm formation (18–20), and flagellar competency is associated with dissemination and pathogenesis (21–24). *P. aeruginosa* displays several well-established forms of motility: swimming, twitching, and swarming. During swimming motility *P. aeruginosa* uses a single polar flagellum to propel through liquid (25), while for twitching, type IV pili extension and retraction are required for movement on hard surfaces (26). Swarming is a social behavior with a characteristic tendril formation on semi-solid surfaces that requires flagella and the secretion of rhamnolipids as a surfactant, while pili also contribute (27, 28). Another type of motility that has been described is ‘surfing’, where *P. aeruginosa* spreads over semi-solid media containing the glycoprotein mucin, that is abundant in the mucus accumulated in the CF airways and gastrointestinal tract (29, 30). This motility is thought to depend on the viscous nature and lubricant properties of mucin and requires *P. aeruginosa* flagellar function while pili and rhamnolipids are expendable (29, 31). *P. aeruginosa* also exhibits sliding motility which requires only the production of rhamnolipids, but no motility appendages (32). Therefore, physical parameters are an important cue for *P. aeruginosa*, where the viscosity of the environment dictates the type of motility, with liquid and low viscosity environments facilitating swimming, and hard surfaces inducing twitching.

*P. aeruginosa* motility is affected by interspecies interactions. Fungal volatiles farnesol and ethanol at high concentrations inhibit *P. aeruginosa* motility, while lower concentrations of ethanol induce swarming when carbon sources are limited, possibly signaling the availability of nutrients (33–35). Further, *P. aeruginosa* is commonly found in concert with *Staphylococcus aureus* in wound infections and nosocomial pneumonia, and these two species are the most frequently isolated bacterial pathogens from the airways of people with CF (36). It has been observed that *P. aeruginosa* engages in directional pili-based ‘exploratory motility’ via the Pil-Chp chemosensory system in response to the *S. aureus* secreted cytotoxins phenol-soluble modulins (PSMs) (37, 38).

In this study, we investigate whether additional *P. aeruginosa* motility behaviors are altered by *S. aureus* exoproducts. We show that the presence of surfactants from *S. aureus*, the soil bacterium *Bacillus subtilis*, and interkingdom species, such as fungi, plants, and humans, as well as synthetic surfactants enables a flagellar-based surface spreading motility in *P. aeruginosa*. Bacterial surface motility has been classified into four types: swarming, twitching, sliding, and gliding (39), and we demonstrate that surface spreading facilitated by exogenous surfactants is an additional distinct type of surface motility. Flagellar-based surface spreading in the presence of exogenous surfactants is not a general bacterial response, but specific to *P. aeruginosa*. This motility occurs on semi-solid as well as hard agar surfaces, where *P. aeruginosa* is otherwise non-motile, thereby demonstrating an emergent phenotype by co-opting already existing interspecies surfactants from the environment. Finally, a comparison with mucin-mediated surfing suggests that this surfactant-facilitated surface spreading is a surfing-like motility, although only movement on mucin requires a quorum sensing regulator.

## RESULTS

### *P. aeruginosa* displays emergent surface spreading in the presence of *S. aureus* secreted products

To determine the effect of *S. aureus* secreted products on *P. aeruginosa* motility, we performed *P. aeruginosa* twitching, swimming, and swarming assays in the presence of 25% (v/v) *S. aureus* cell-free supernatant, or media salts as a control. As reported previously (37), we observed that *P. aeruginosa* twitched farther with the addition of *S. aureus* supernatant **(Supp. Fig. 1A)**. In the swimming assay, while *P. aeruginosa* swam uniformly creating a circular swim area in the control agar, the swim patterns were irregular and oval-shaped in the presence of *S. aureus* supernatant **(Supp. Fig. 1B)**.

In a standard swarm assay on semi-solid agar plates (containing 0.5% agar) *P. aeruginosa* swarmed with characteristic tendril formation on the control plate (**Fig. 1A**). However, on plates with 25% *S. aureus* supernatant, *P. aeruginosa* exhibited a motility unlike swarming where the bacteria spread out across the plate, and then showed some swarming-like tendril formation (**Fig. 1A**). *P. aeruginosa* exhibited similar motility on agar containing 12.5% *S. aureus* supernatant and an intermediate motility on 5% supernatant, while swarming resumed on plates with 1% supernatant (**Fig. 1A**). Next, we tested if this phenotype occurs on hard agar plates (containing 1.5% agar) on which *P. aeruginosa* is traditionally unable to move. While *P. aeruginosa* did not move on the hard agar control or on the plates containing 5% and 1% *S. aureus* supernatant (**Fig. 1B**), it exhibited emergent surface spreading on the plates containing 25% and 12.5% *S. aureus* supernatant (**Fig. 1B**). There was no tendril formation on hard agar, likely because the tendrils are due to swarming after surface spreading. Thus, *S. aureus* secreted products enable a distinct form of *P. aeruginosa* motility, even in conditions where *P. aeruginosa* is normally nonmotile.

**Figure 1.**
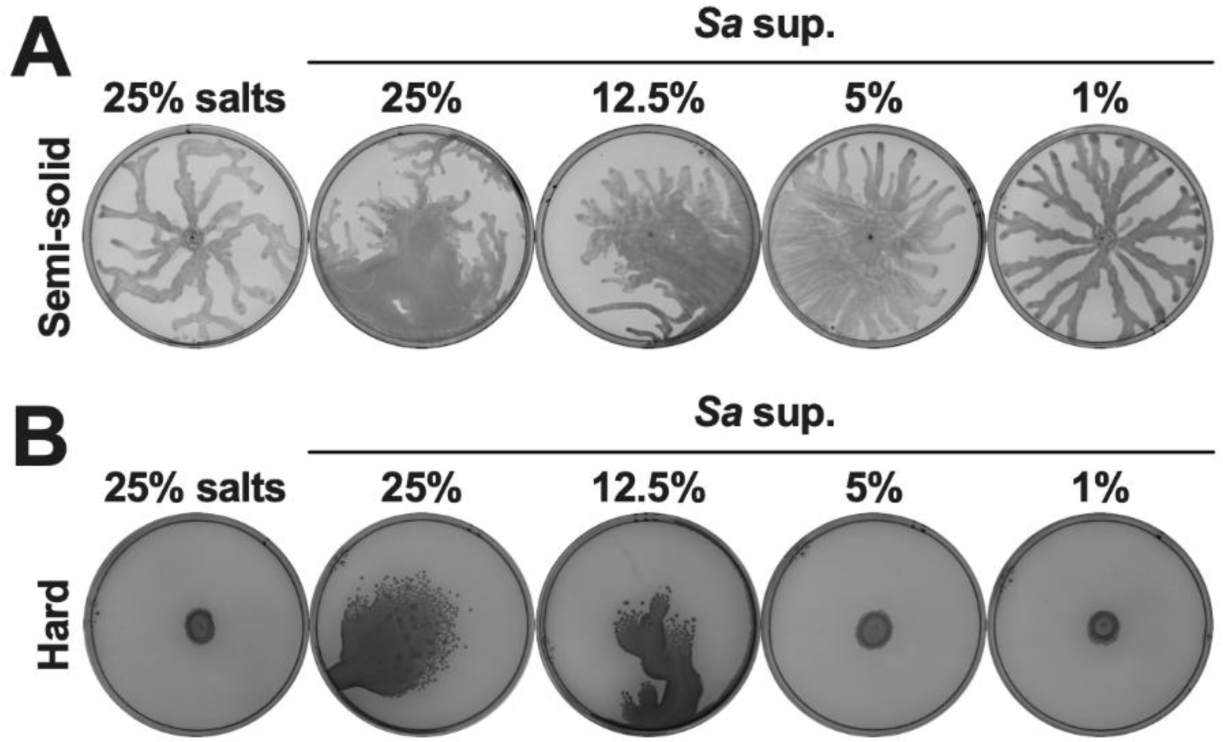
*S. aureus* secreted products enable *P. aeruginosa* surface spreading in a dose-dependent manner. *P. aeruginosa* was inoculated on **(A)** semi-solid or **(B)** hard agar plates containing the indicated percentages of media salts as a control or *S. aureus* supernatant, and plates were imaged after 24 hours incubation. **(A,B)** Representative images of three independent replicates are shown. Additional replicates are shown in **Extended** Figure 1.

To determine the generality of this motility, we performed the surface spreading assay with clinical isolates of both species. We tested four clinical isolates of *P. aeruginosa* collected from different people with CF (CF017, CF033, CF057, and CF095) **(Supp. Table 1**) on both semi-solid and hard agar and observed that the isolates CF017 and CF057 swarmed and exhibited surface spreading to varying degrees, while isolates CF033 and CF095 did not **(Supp. Fig. 2A and B**). To determine if these isolates also had other motility deficiencies, we tested these isolates for swimming and twitching. Isolate CF017 swam similar to *P. aeruginosa* PA14, while isolates CF033, CF057, CF095 all had swimming deficiencies **(Supp. Fig. 3A**). Additionally, the twitch zone of CF033 was similar to PA14, but CF017 and CF057 had smaller twitch zones and CF095 did not twitch **(Supp. Fig. 3B**). Therefore, the isolates that do not exhibit the emergent motility may also have flagellar and/or pili deficiencies. These *P. aeruginosa* strains were co-isolated with four *S. aureus* strains, CF019, CF032, CF058, and CF100, respectively **(Supp. Table 1)**. Cell-free supernatants from each of these *S. aureus* clinical isolates could facilitate this motility in *P. aeruginosa* PA14 **(Supp. Fig. 4A and B)**. These results demonstrate that while there is some variation, clinical strains of *P. aeruginosa* and *S. aureus* can exhibit and enable surface spreading, respectively.

### Interspecies surfactants facilitate surface spreading in *P. aeruginosa*

To identify the molecules that enable surface spreading in *P. aeruginosa*, we investigated whether other species promote this motility, and found that *P. aeruginosa* also exhibited similar motility on plates with supernatant from *P. aeruginosa* and *Bacillus subtilis* ZK3814 (**Fig. 2A**) but swarmed on plates with supernatant from *B. subtilis* PY79, *Klebsiella pneumoniae, Vibrio cholerae, Escherichia coli, Burkholderia cenocepacia* BAA245, and *B. cenocepacia* 25608 (**Fig. 2A** and **B**).

**Figure 2.**
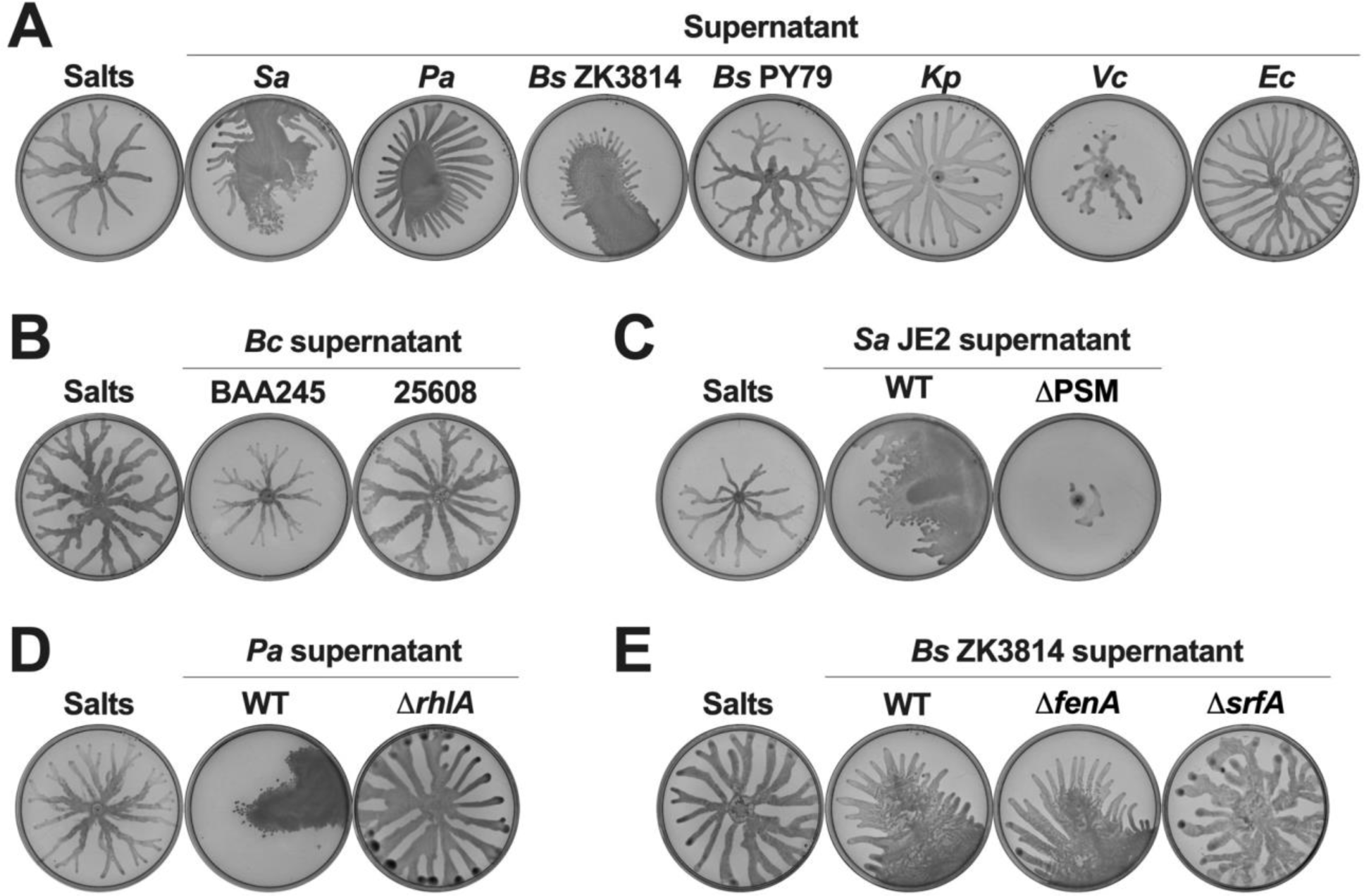
Interspecies secreted surfactants facilitate surface spreading in *P. aeruginosa*. **(A-E)** *P. aeruginosa* was inoculated on semi-solid agar plates containing 25% media salts as a control or supernatant from the indicated species: *S. aureus* (JE2), *P. aeruginosa* (PA14), *B. subtilis, K. pneumoniae, V. cholerae, E. coli* (MG1655), and *B. cenocepacia*. Supernatant was from **(A, B)** WT strains and **(C-E)** mutants for the biosynthesis of surfactants. Images were taken after 24 hours incubation. Representative images of three independent replicates are shown. Additional replicates are shown in **Extended** Figure 2.

A common class of molecules previously implicated in motility, and known to be secreted by each of the inducing strains, but none of the non-inducing strains, is biosurfactants (40). *S. aureus* produces phenol soluble modulins (PSMs) (41–43), *P. aeruginosa* produces rhamnolipids (44), and *B. subtilis* ZK3814 produces fengycin and surfactin (45). We therefore tested the supernatant of mutants deficient in each of these biosurfactants for their ability to facilitate motility in *P. aeruginosa*. The supernatants of the *S. aureus* JE2 and LAC strains that lack production of PSMs did not enable motility in *P. aeruginosa* (**Fig. 2C** and **Supp. Fig. 5A, B, and C)**. Further, supernatant from *agrA*::tn, *agrB*::tn, or *agrC*::tn, mutants of the *S. aureus agr* quorum sensing system which is required for the production of PSMs, also did not lead to *P. aeruginosa* surface spreading **(Supp. Fig. 5A and D**). Similarly, we found that *P. aeruginosa* did not exhibit this motility on plates containing supernatant from the *P. aeruginosa* Δ*rhlA* mutant deficient for rhamnolipids (**Fig. 2D** and **Supp. Fig. 6A**). Finally, supernatant from a *B. subtilis* mutant without surfactin production (Δ*srfA*) was unable to promote *P. aeruginosa* surface spreading, unlike the parental WT *B. subtilis* ZK3814 as well as Δ*fenA*, a mutant deficient for fengycin production (**Fig. 2E** and **Supp. Fig. 6B**). Collectively, these data show that the production of biosurfactants is required for the interspecies facilitated surface spreading in *P. aeruginosa*.

### Detergents and surfactants are sufficient to facilitate surface spreading in *P. aeruginosa*

Given that bacterial-secreted surfactants were necessary to facilitate surface spreading in *P. aeruginosa,* we investigated if the addition of biological and synthetic surfactants was sufficient to enable this motility. We tested commercially available biosurfactants from the motility-inducing species, including rhamnolipids from *P. aeruginosa* and surfactin from *B. subtilis;* mucin, which has been previously implicated in *P. aeruginosa* surfing motility (31); and the synthetic surfactants sodium dodecyl sulfate (SDS), cetrimonium bromide (CTAB), and Triton X-100. *P. aeruginosa* exhibited spreading motility on plates containing all these surfactants to varying degrees, except CTAB, which inhibited motility (**Fig. 3A**). Further, we tested biosurfactants from interkingdom species, such as mannosylerythritol lipid A (MEL-A) and sophorolipid from yeasts, plant-produced saponin, and human cathelicidin LL-37 and pulmonary surfactant dipalmitoylphosphatidylcholine (DPPC). All these interkingdom surfactants, except DPPC, caused the spreading motility, especially on semi-solid agar (**Fig. 3B**). Thus, diverse biological and synthetic surfactants are sufficient to enable *P. aeruginosa* surface spreading motility.

**Figure 3.**
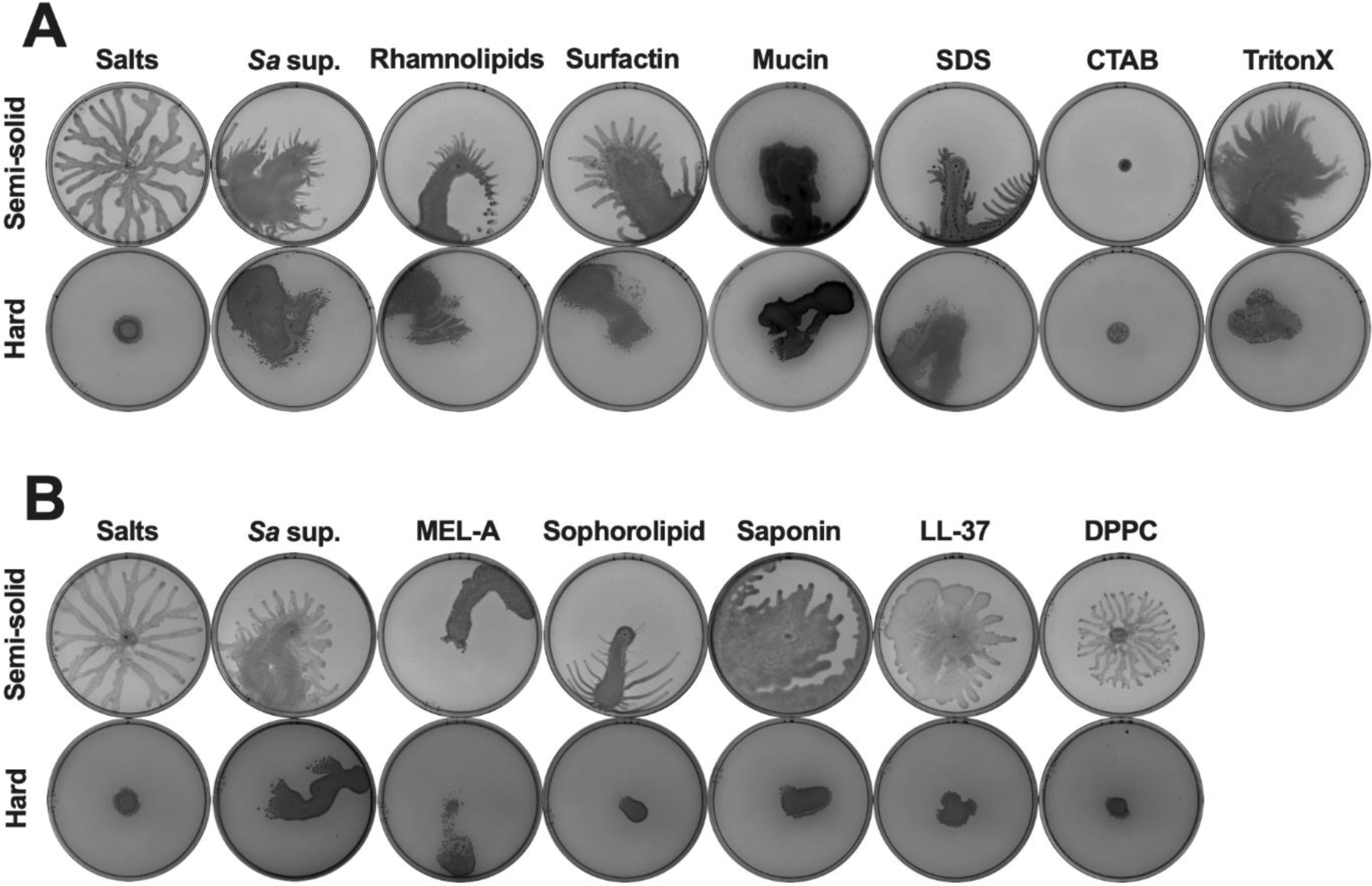
The addition of diverse biotic and synthetic surfactants is sufficient to enable surface spreading in *P. aeruginosa*. **(A,B)** *P. aeruginosa* was inoculated on (top) semi-solid or (bottom) hard agar plates containing **(A)** (from left to right) 25% media salts as a control or *S. aureus* supernatant, or 25% media salts with the addition of 50 µg·mL^-1^ rhamnolipids, 5 µg·mL^-1^ surfactin, 0.4% mucin, 0.1% SDS, 0.1% CTAB, or 0.1% Triton X-100; or **(B)** 25% media salts control or *S. aureus* supernatant; or 25% media salts with the addition of 25 µg·mL^-1^ MEL-A, 25 µg·mL^-1^ sophorolipid, 25 µg·mL^-1^ saponin, 100 µg·mL^-1^ LL-37, or 100 µg·mL^-1^ DPPC. **(A,B)** Images were taken after 24 hours incubation. Representative images of three independent replicates are shown. Additional replicates are shown in **Extended** Figure 3.

### *P. aeruginosa* surface spreading requires flagella, but not pili or rhamnolipids

*P. aeruginosa* swarming motility requires endogenous production of the biosurfactant rhamnolipids, therefore we tested whether surface spreading was similar to swarming, by comparing their respective genetic requirements. Pili, flagella, and the surfactant rhamnolipids all contribute to *P. aeruginosa* swarming (27, 28). Additionally, swarming is also affected by quorum sensing systems and other regulatory pathways (27, 46, 47).

First, we tested several mutants that lack the following structural components of the *P. aeruginosa* flagella: the proximal rod protein FlgB, the hook protein FlgE, or the flagellin subunit FliC (48), and are therefore defective in swimming **(Supp. Fig. 7A**) and swarming (**Fig. 4A**). On the semi-solid and hard agar plates with *S. aureus* supernatant, the *flgB::*tn, *flgE::*tn, and Δ*fliC* mutants only slid on the plates without spreading out, unlike the motility exhibited by the WT strain (**Fig. 4A** and **Supp. Fig. 7B**), indicating that flagella are required for surface spreading, and that sliding is likely passive, and occurs only in the presence of surfactants possibly due to a slight difference in the leveling of the agar. To examine the motility in a species that does not encode flagella, we tested *S. aureus* itself, and found that it was immobile on the control agar and slid on the plates with *S. aureus* supernatant like the *P. aeruginosa* flagellar mutants (**Fig. 4B** and **Supp. Fig. 7C**). To determine if these flagella-deficient strains could move in the presence of motile *P. aeruginosa*, these strains were fluorescently labeled, mixed, and added to hard agar with *S. aureus* supernatant. As expected, two distinctly labeled *P. aeruginosa* WT strains co-cultured on hard agar exhibited surface spreading equally (**Fig. 4C**). However, the Δ*fliC* mutant and *S. aureus* still only slid on the plate even in co-culture with *P. aeruginosa* WT cells that showed surface spreading (**Fig. 4C**), suggesting that this motility is an active process, unlike passive sliding, and the presence of a motile population cannot compensate for loss of competent flagella. Hence, *P. aeruginosa* surface spreading in the presence of interspecies surfactants requires active flagellar function.

**Figure 4.**
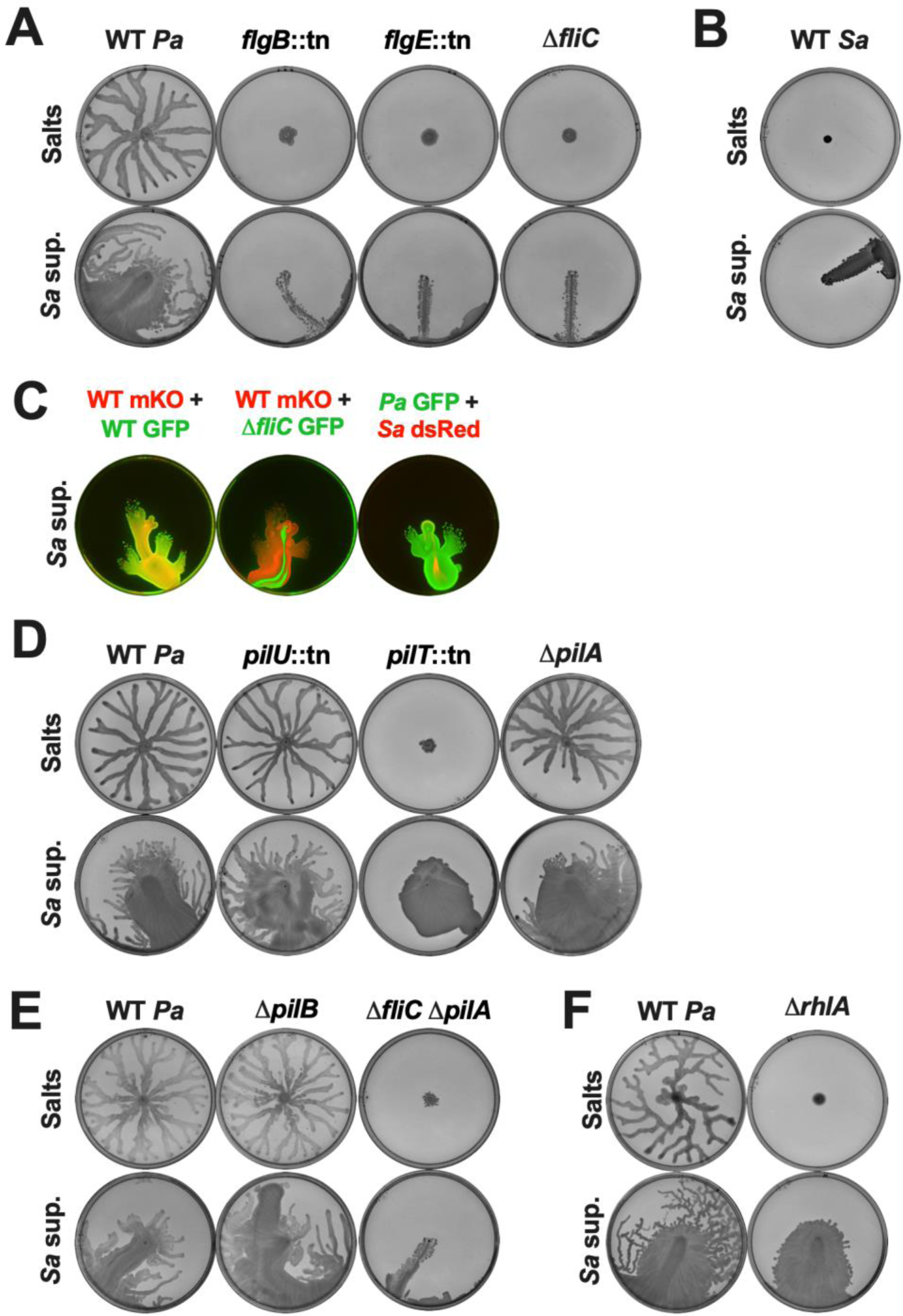
Surface spreading requires flagella, but not pili or rhamnolipids. The indicated strains of **(A,D,E,F)** *P. aeruginosa* or **(B)** *S. aureus* were inoculated on **(A,B,D,E,F)** semi-solid agar plates containing 25% media salts as a control or *S. aureus* supernatant as indicated. **(C)** From left to right, *P. aeruginosa* WT-mKO mixed with WT-GFP, *P. aeruginosa* WT-mKO mixed with Δ*fliC-*GFP, and *P. aeruginosa* WT-GFP mixed with *S. aureus* WT*-*dsRed were inoculated on hard agar plates containing 25% *S. aureus* supernatant. **(A-F)** Motility was imaged after 24 hours incubation. Representative images of three independent replicates are shown. Additional replicates are shown in **Extended** Figure 4.

We next determined if pili, which are required for twitching and contribute to swarming (27), are necessary for this motility. We tested multiple twitching-deficient pili mutants **(Supp. Fig. 8A and B)** - two mutants deficient in pili retraction, *pilT*::tn and *pilU*::tn (49), two mutants lacking the extended pili fiber, Δ*pilB*, via deficiency in pili extension (50), and Δ*pilA* due to the lack of the pilin subunit (26), and two mutants deficient for pili-mediated chemotaxis, Δ*pilJ* and Δ*pilG* (51). While these mutants showed differences in swarming, all of these strains, displayed surface spreading on plates with *S. aureus* supernatant (**Fig. 4D and E**, **Supp. Fig. 8C and D, Supp. Fig. 9A**). In particular, the *pilT*::tn mutant was unable to swarm, but still showed surface spreading, albeit without tendril formation, suggesting that the surface spreading is distinct from swarming, although the tendril formation likely indicates swarming.

Previously, it was demonstrated that *P. aeruginosa* shows passive ‘sliding motility’ that does not require any appendages but is impeded by pili (32). In that study, flagellar mutants did not slide, while pili mutants and a mutant for both pili and flagella showed sliding motility (32). We therefore tested a Δ*fliC* Δ*pilA* double mutant and found that this mutant only slid on the plates with *S. aureus* supernatant and did not demonstrate surface spreading (**Fig. 4E**). Taken together, *P. aeruginosa* surface spreading in the presence of *S. aureus* secreted products is an active motility that requires flagella but not pili.

While swarming, *P. aeruginosa* secretes rhamnolipids, biosurfactants that lubricate the area and allow the cells to move over the surface (27, 28). We found that while a mutant deficient in endogenous rhamnolipid production, Δ*rhlA*, was unable to swarm as previously reported (27, 28), it still spread on both semi-solid and hard agar plates containing *S. aureus* cell-free supernatant (**Fig. 4F** and **Supp. Fig. 9B**), suggesting that endogenously produced rhamnolipids are not required for surface spreading. This mutant did not exhibit tendril formation after the surface spreading on semi-solid agar, reiterating that the tendrils likely indicate swarming.

Finally, quorum sensing and other motility regulators are known to be important for swarming (27, 46, 47). *P. aeruginosa* utilizes three quorum sensing systems: Las, Rhl, and PQS (52). To examine the role of quorum sensing in surface spreading we tested mutant strains lacking each individually (Δ*rhlR,* Δ*lasR,* Δ*pqsA*), or in combination (Δ*rhlR* Δ*lasR* Δ*pqsA*). The Δ*rhlR,* Δ*lasR,* and Δ*rhlR* Δ*lasR* Δ*pqsA* mutants did not swarm on the control plates, while the Δ*pqsA* mutant did **(Supp. Fig. 10A**). However, all mutants displayed surface spreading on the plates with *S. aureus* supernatant to varying degrees, with the Δ*pqsA* strain being hyper-motile compared to the other strains and covering the entire semi-solid agar plate **(Supp. Fig. 10A and B)**. We also tested mutants from additional regulatory pathways that control motility, including the Gac/Rsm and Wsp pathways (53–56). The Δ*retS* and Δ*rsmA* mutants did not swarm, while the Δ*gacA,* Δ*wspR,* and Δ*toxR* mutants exhibited swarming **(Supp. Fig. 11)**. However, all these regulatory mutants exhibited surface spreading **(Supp. Fig. 11)**, indicating that these regulatory pathways are not required for *P. aeruginosa* surface spreading, although quorum sensing may affect motility patterns. Thus, this interspecies surfactant-facilitated motility is distinct from swarming since it does not require processes that are necessary for swarming, such as pili retraction or quorum sensing and other regulatory pathways.

### Other bacterial species show distinct motility phenotypes in the presence of exogenous surfactants

Given that *P. aeruginosa* requires flagella for surface spreading, and the non-motile *S. aureus* only slides in the presence of exogenous surfactants, we tested other bacterial species like *B. cenocepacia*, *K. pneumoniae*, *V. cholerae*, and *E. coli* on semi-solid plates containing *S. aureus* supernatant, Triton X-100, or saponin **(Supp. Fig. 12)**. All these species, except *K. pneumoniae*, are known to be flagellated (57, 58), and *E. coli* and *B. cenocepacia* demonstrated swimming motility (**Supp Fig. 13A and B**). While *V. cholerae* showed some surface spreading in the presence of *S. aureus* supernatant, none of the other species showed robust movement beyond sliding. Thus, surface spreading as exhibited by *P. aeruginosa* is not a ubiquitous bacterial motility, even for flagellated swimming-proficient species in the presence of exogenous surfactants.

We also tested *B. subtilis* strains ZK3814 that demonstrated swimming motility (**Supp Fig. 13A**) and produces surfactin and PY79 that does not, on semi-solid agar containing either media salts as a control, or *S. aureus* supernatant, rhamnolipids, or saponin **(Supp. Fig. 14)**. Interestingly, *B. subtilis* ZK3814 swarmed in all these conditions, and did not switch to surface spreading, unlike what is seen in *P. aeruginosa*. Further, *B. subtilis* PY79 was nonmotile on the control plate, but showed similar motility to *B. subtilis* ZK3814 in the presence of exogenous surfactants, possibly preceded by sliding and surface spreading. Therefore, the presence of exogenous surfactants likely restores swarming motility in surfactant-deficient *B. subtilis*, unlike the *P. aeruginosa* rhamnolipid-deficient mutant, which displayed only surface spreading (**Fig. 4F**).

### Cells undergoing surface spreading do not differentially transcriptionally regulate motility pathways

Given the lack of requirement of all major motility regulators for surface spreading, and the diversity of surfactant molecules that facilitate it, this motility could potentially be regulated by alternative transcriptional regulators or exclusively by the physical cue of reduced surface tension. Thus, to test for transcriptional regulation during surface spreading, we performed RNA sequencing (RNA-seq) on *P. aeruginosa* cells scraped from the leading edge on semi-solid or hard agar plates containing the media salts control or *S. aureus* supernatant at 17 hours **(Supp. Fig. 15A)**. We found approximately 100 to 350 genes that were upregulated and downregulated on the supernatant compared to the media salts control in both cases **(Supp. Tables 2 and 3)**. We reasoned that the differentially regulated genes would include genes specific to this motility as well as genes that were regulated upon exposure to unrelated *S. aureus* secreted molecules. To distinguish between generalized responses to *S. aureus* products and motility specific transcripts, we performed a secondary comparison of both motility conditions to genes we previously identified as differentially regulated in early log-phase *P. aeruginosa* planktonic cultures 2 hours after the addition of *S. aureus* supernatant compared to media salts as a control (59), and found significant overlaps (**Fig. 5A**). Some of the highest upregulated genes in both motility conditions as well as in planktonic conditions were genes involved in tricarboxylic acids (TCA) uptake and acetoin catabolism, likely due to *S. aureus* secreted citrate and acetoin, respectively (59) (**Fig. 5B**, **Supp. Fig. 15B and C, Supp. Tables 3, 4, and 5)**. Next, we performed Gene Ontology (GO) enrichment analysis of the *P. aeruginosa* genes differentially regulated in the individual motility conditions, both motility conditions (semi-solid and hard agar plates), as well as those regulated exclusively in both motility conditions, but not in the planktonic phase (60–62) (**Fig. 5C**, **Supp. Fig. 15D and E, and Supp. Tables 4, 5, and 6)**. Genes involved in aerobic respiration and response to heat were enriched in the genes downregulated in both motility conditions **(Supp. Fig. 15E and Supp. Table 6)**, but these genes were also downregulated in the planktonic phase and are likely unrelated to the motility **(Supp. Fig. 15E and Supp. Tables 5 and 6)**.

**Figure 5.**
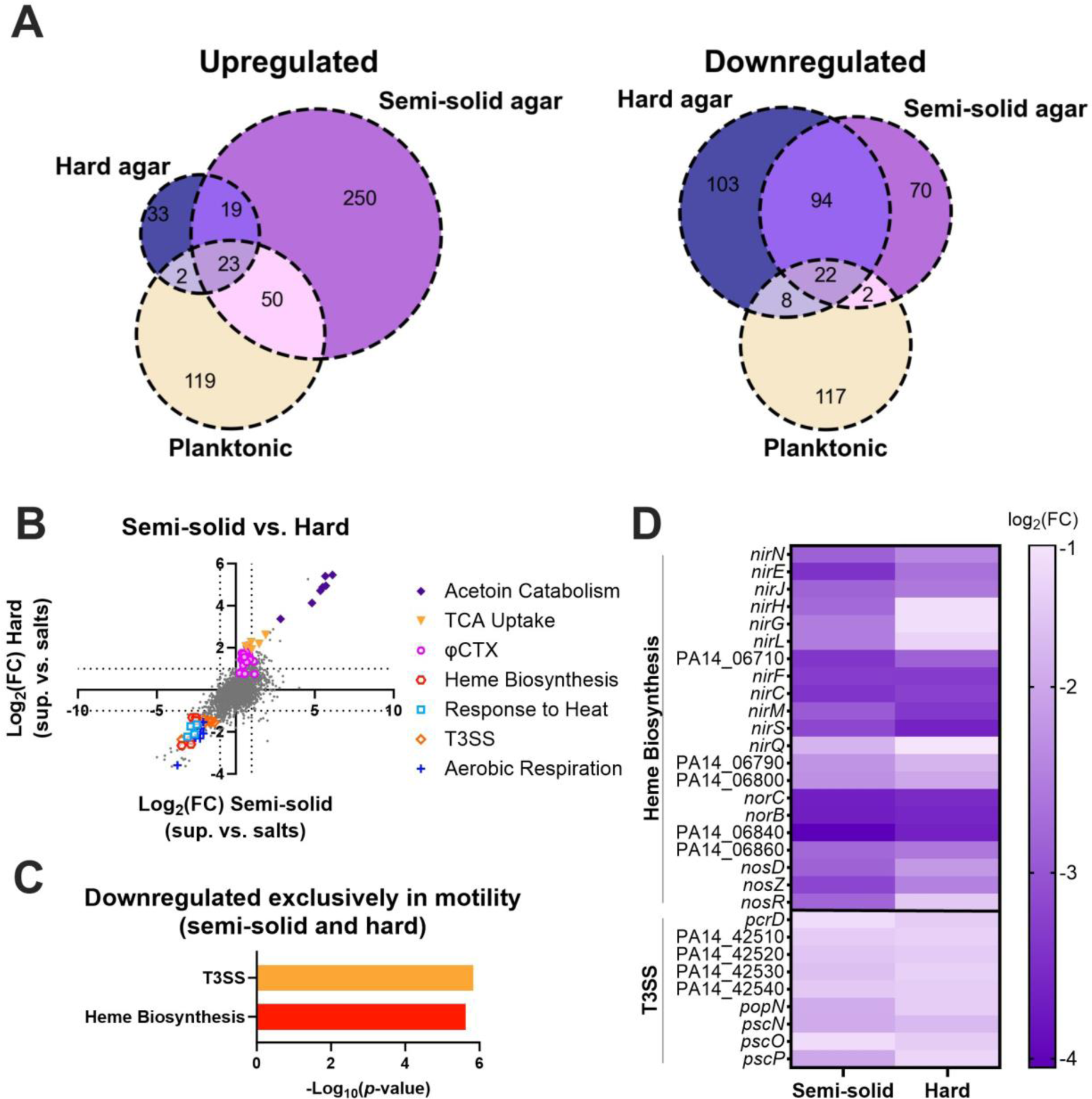
*P. aeruginosa* differentially regulates genes involved in T3SS, and heme biosynthesis, but not motility, during surface spreading on *S. aureus* secreted products. Transcript levels were compared between *P. aeruginosa* cells exposed to *S. aureus* supernatant and those exposed to media salts as a control on semi-solid and hard agar after 17 hours. **(A)** Venn diagrams of upregulated and downregulated genes on hard and semi-solid agar, as well as in planktonic *P. aeruginosa* cells after *S. aureus* supernatant exposure compared to media salts control (59). **(B)** Scatter plot of log_2_(fold-change) transcript levels on hard agar compared to the log_2_(fold-change) of transcript levels on semi-solid agar. Differentially regulated genes in pathways enriched in the common downregulated genes, as well as in TCA uptake, acetoin catabolism, and φCTX are shown. **(C)** GO enrichment of *P. aeruginosa* genes differentially expressed in response to *S. aureus* supernatant on both semi-solid and hard agar plates, but not in planktonic phase (60–62). Nonredundant categories are shown. **(D)** Heatmap showing the log_2_(fold-change) of genes associated with enriched pathways for heme biosynthesis and T3SS on semi-solid and hard agar.

The *P. aeruginosa* genes differentially regulated exclusively in both motility conditions, but not in the planktonic phase, were not enriched for any motility pathways (**Fig. 5C** and **Supp. Table 6)**, suggesting that transcriptional regulation may not play an important role during surface spreading, consistent with our results that major motility regulators are not required for this motility **(Supp. Fig. 10 and 11)**. Instead, genes downregulated exclusively in the motility conditions showed enrichment of the Type 3 Secretion System (T3SS) and heme biosynthesis pathways, while the upregulated genes showed no statistically enriched pathways (**Fig. 5C** and **D**, **and Supp. Table 6**). Additionally, on the hard agar there was an upregulation in transcripts from the phage φCTX that encode the R and F-type pyocins (**Fig. 5B**, **Supp. Fig. 15B, and Supp. Table 3**)(63), indicating pyocin induction in this condition. This likely explains the presence of plaque-like clearings on the hard agar (e.g. in **Fig. 1B**), which was especially apparent for the LasR mutant **(Supp. Fig. 10**). Multiple cellular stresses can lead to both induction of prophage and downregulation of T3SS (64–67), indicating that *P. aeruginosa* senses physiological stress while undergoing surface spreading. Further, heme biosynthesis and associated denitrification genes (e.g. *nirS*, *nirQ*, *norC, norB)* are induced in anaerobic conditions (68). The relative downregulation of these genes during surface spreading may be indicative that the control cells (swarming cells in the semi-solid agar and non-motile cells in the hard agar) are switching to anaerobic metabolism, while cells undergoing surface spreading continue with aerobic respiration. These results indicate that *P. aeruginosa* surface spreading motility is unlikely to be regulated at the transcriptional level, although cells may experience physiological stress and metabolic changes while undergoing surface spreading in the presence of *S. aureus* exoproducts.

### *P. aeruginosa* surfactant-mediated surface spreading is a surfing-like motility

We next tested whether interspecies surfactant-facilitated surface spreading is similar to mucin-induced surfing (29, 31, 69, 70), by comparing the genetic requirements for surface spreading on semi-solid plates containing *S. aureus* supernatant or mucin. Flagellar and pili mutants showed similar phenotypes for motility on each, where *flgB*::tn, and Δ*fliC* showed no motility, while *pilT*::tn and Δ*pilA* moved on both **(Supp. Fig. 16A)**.

Previously, Rhl and Las as well as components of Gac/Rsm were reported to contribute to mucin-based surfing (29, 69, 70). We observed that while the Δ*pqsA* mutant moved on both the *S. aureus* and mucin plates, the Δ*rhlR* and Δ*rhlR* Δ*lasR* Δ*pqsA* mutants moved moderately on *S. aureus* supernatant and had no movement on the plates with mucin **(Supp. Fig. 16B)**. Interestingly, the Δ*lasR* mutant moved on the *S. aureus* supernatant but not on mucin, demonstrating a divergence of quorum sensing requirements for mucin-based surfing (29, 31) and surfactant-enabled motility. The Δ*retS*, Δ*gacA*, Δ*rsmA*, Δ*wspR*, and Δ*toxR* regulatory mutants all moved on both *S. aureus* supernatant and mucin to varying degrees **(Supp. Fig. 17)**. To further delineate the role of LasR in motility on a variety of surfactants, we tested the Δ*lasR* mutant on semi-solid plates with supernatants from *S. aureus*, *P. aeruginosa*, and *B. subtilis* ZK3814, as well as with the addition of mucin and the surfactant Triton X-100. While the Δ*lasR* mutant did not move on mucin, it exhibited surface spreading on all of the other surfactant-containing plates (**Fig. 6A**). Overall, the surfactant-dependent surface spreading we observed, and mucin-dependent surfing motility have similar appendage and regulatory requirements, indicating that surfactant-based surface spreading is a surfing-like motility.

**Figure 6.**
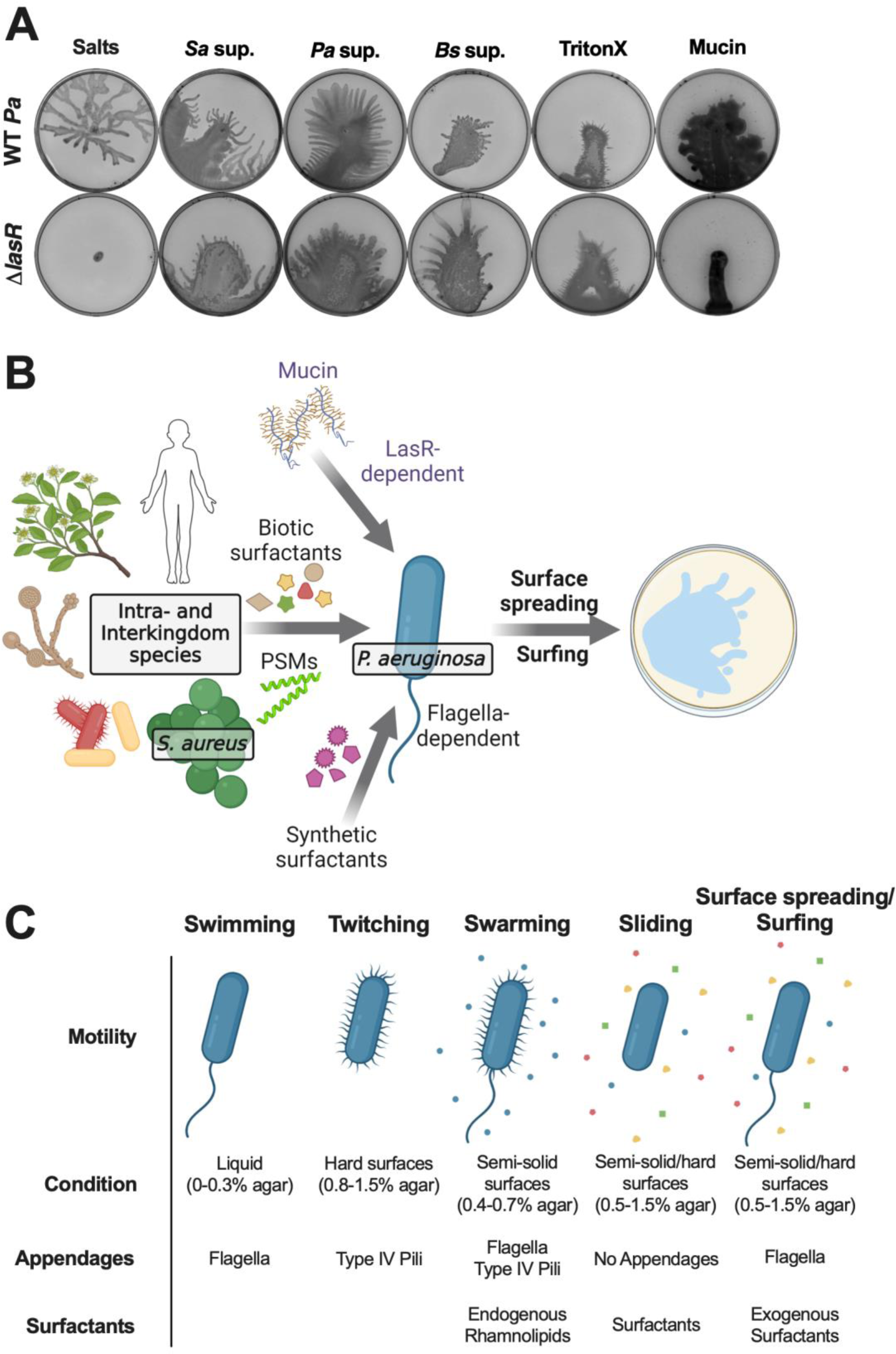
The Las system is required for surfing motility on mucin, but not on surfactants. **(A)** *P. aeruginosa* WT and Δ*lasR* were inoculated on semi-solid agar plates containing (from left to right) 25% media salts as a control or supernatant from the indicated species: *S. aureus* (JE2), *P. aeruginosa* (PA14), or *B. subtilis* (ZK3814); or 25% media salts with the addition of 0.1% Triton X-100 or 0.4% mucin. Images were taken after 24 hours incubation. Representative images of three independent replicates are shown. Additional replicates are in **Extended** Figure 6. **(B)** Working model of *P. aeruginosa* motility in response to exogenous surfactants. The glycopeptide mucin, PSMs secreted from *S. aureus*, and other biotic surfactants secreted by human cells, plants, yeast, and bacteria, as well as synthetic surfactants, are co-opted by *P. aeruginosa.* In the presence of exogenous surfactants, *P. aeruginosa* exhibits flagella-dependent spreading motility on surfaces. In contrast to surfactant-facilitated surface spreading, mucin-mediated surfing motility requires the LasR quorum sensing system in *P. aeruginosa*. **(C)** (Left to right) *P. aeruginosa* swimming motility occurs in liquid conditions and requires flagella (25), twitching motility occurs on hard surfaces and requires type IV pili (26), swarming motility occurs on semi-solid surfaces and requires flagella, type IV pili, and rhamnolipids (27, 28), sliding motility occurs on semi-solid surfaces and endogenous or exogenous surfactants (32), and surface spreading/surfing motility occurs on semi-solid and hard surfaces and requires flagella and exogenous surfactants (29, 31, 69, 70).

## DISCUSSION

Bacterial locomotion is an advantageous trait that enables migration to more favorable environments and access to fresh nutrients. Within microbial communities, interspecies secreted factors can affect bacterial motility, by activation (8), inhibition (9), or endowing nonmotile strains the capability to move (13, 14). Many of these observations were reported by examining pairwise interactions, which can reveal novel behavioral phenomena as well as the underlying mechanisms (11). Here, we interrogated how *P. aeruginosa* motility is affected by *S. aureus* secreted molecules and uncovered a surfing-like surface spreading motility that depends on the presence of exogenous interspecies or synthetic surfactants (**Fig. 6B**), likely due to the low surface tension environment provided by the physical properties of these surfactants. The surfactant-based motility we observed requires flagellar function, but not type IV pili or endogenous production of rhamnolipids, suggesting it is distinct from previously described motility types in *P. aeruginosa* (**Fig. 6C**). It permits the emergence of motility when *P. aeruginosa* is typically nonmotile and is facilitated by surfactants from a variety of species, including co-infecting pathogens, putative environmental neighbors, as well as infection hosts, demonstrating that *P. aeruginosa* can take advantage of pre-existing public goods from nearby species to explore new niches.

We observed that in the presence of exogenous surfactants, *P. aeruginosa* cells spread over the surface, an active process that required flagellar function. This was followed by swarming tendril formation on semi-solid agar, or likely dispersal of individual cells on hard agar that resulted in colonies at the mobile leading edge. The motility was facilitated by the presence of diverse types of biological and synthetic molecules of a variety of chemical compositions, suggesting that their shared properties of reducing surface tension and acting as wetting agents led to the motility. Previously it had been shown that when other gelling agents, the polysaccharides gellan gum and carrageenan, are used in place of agar in semi-solid media, *P. aeruginosa* exhibits a similar flagellar-dependent, rhamnolipid-independent surface spreading motility, possibly because the alternate gelling agents also acted as wetting agents (71). It is possible that the *P. aeruginosa* polar flagellum directly senses surfactant properties by the alleviation of impediments on flagellar rotation (72), and transitions cells to surface spreading.

The reduced surface tension due to exogenous surfactants likely allows for increased movement, resulting in the emergence of motility even on hard agar, where *P. aeruginosa* is otherwise nonmotile, and enabling surface motility without rhamnolipid production, unlike what is seen for swarming. Intrinsic rhamnolipid production occurs and is required during swarming, while the presence of exogenous rhamnolipids leads to the emergent motility phenotype (**Fig. 3A**). Given that *P. aeruginosa* rhamnolipid production is quorum-sensing dependent and requires high cell density, it is likely that in the presence of exogenous surfactants, *P. aeruginosa* cells rapidly sense the change in environmental conditions resulting in surface spreading motility before they produce rhamnolipids.

While reduced surface tension likely facilitates *P. aeruginosa* surface spreading via flagellar function, this is not a general phenotype for bacteria. We observe that *E. coli* and *B. cenocepacia* strains with functional flagella do not show surface spreading, suggesting that additional parameters likely affect this motility. It is possible that cell surface or other physical characteristics of *E. coli* and *B. cenocepacia* prevent surface spreading, unlike what is seen in *P. aeruginosa*. Further, whereas *P. aeruginosa* switches from swarming to surface spreading in the presence of exogenous surfactants, *B. subtilis* swarming is unaltered. Additionally, a *B. subtilis* strain deficient for surfactin production (PY79) resumes swarming in the presence of exogenous surfactants, unlike what is seen for the *P. aeruginosa* rhamnolipid mutant Δ*rhlA* (**Fig. 4F**), which demonstrates only surface spreading. Thus, the *P. aeruginosa* surface spreading in the presence of exogenous surfactants is a motility response distinct from that displayed by other motile bacteria.

We systematically examined the requirement of several regulators that control motility (73), such as components of Pil-Chp, quorum sensing, Gac/Rsm, and Wsp surface sensing, but none were needed for surfactant-based surface spreading, except for mucin-based surfing requiring the LasR quorum sensing regulator (**Fig. 6A**) (29, 69). Nonetheless, there were noticeable differences in the appearance of surface spreading quorum sensing mutants. On semi-solid agar, the final step of tendril formation was absent in Rhl mutants that are expected to lack rhamnolipid production, indicating that the tendrils here are swarming, and require intrinsic production of rhamnolipids. In addition, mutants devoid of the PQS pathway, which plays a role in swarming tendril repulsion (74), are hyper-motile on semi-solid agar, suggesting that the PQS pathway may play a role in limiting surfactant-based surface spreading.

*P. aeruginosa* and *S. aureus* often co-infect wounds and the airways of people with CF. We observed that the presence of *S. aureus* PSMs, which are amphipathic alpha-helical peptides with surfactant properties, facilitates surface spreading motility in *P. aeruginosa*. Previously, these *S. aureus* secreted toxins have been implicated in enabling motility in both species. In *S. aureus*, PSMs allow nonmotile *S. aureus* to slide on surfaces (42). Additionally, *P. aeruginosa* senses PSMs via the Pil-Chp chemosensory system to induce pili-dependent directional ‘exploratory motility’ (37, 75), which is a motility distinct from the one we describe. *P. aeruginosa* explorers require pili and components of the Pil-Chp system, such as the chemoreceptor PilJ (75), which are not necessary for flagellar-dependent surfactant-based surface motility. Sensing of surface hardness and viscosity plays a role in stimulating motility behaviors (76, 77), with twitching and swarming occurring on hard and semi-solid surfaces, respectively (**Fig. 6C**). However, we observed that even on hard agar surface spreading still solely required flagella and not pili, in contrast to the directionality of exploratory motility detected on hard agar (37, 75). PSMs produced by neighboring *S. aureus* cells are also thought to alter the flow of *P. aeruginosa* rhamnolipids, leading to repulsion of *P. aeruginosa* swarm tendrils (78). Thus, there is an overarching role for PSMs in inducing and altering motility in *P. aeruginosa*, and the mode of locomotive response likely differs based on additional environmental stimuli and/or how these molecules are sensed by *P. aeruginosa*.

In addition to *S. aureus*-produced PSMs (42), many microbes secrete diverse classes of biosurfactants that enable locomotion (79), such as the lipopeptide surfactin by *B. subtilis* (80), the glycolipids rhamnolipids by *P. aeruginosa* and *B. cenocepacia* (81, 82), and the aminolipid serrawettin by *Serratia* spp. (83). As secreted factors, interspecies surfactants can serve as public goods and enable other species to co-opt these wetting agents for motility, and we observed that *P. aeruginosa* can utilize diverse intra-and interkingdom surfactants for this surfing-like motility. It has been observed that in conditions where *B. cenocepacia* is individually non-motile (and does not produce surfactant), it can use *P. aeruginosa* rhamnolipids for motility (84, 85). Further, flagella-deficient *P. aeruginosa* that produces rhamnolipids is incapable of swarming but can co-swarm with *B. cenocepacia* that is using the secreted surfactant (85), providing benefits for each species. Despite this display of hitchhiking on motile *B. cenocepacia*, in our study we did not observe cooperation or hitchhiking between surface spreading *P. aeruginosa* and non-spreading flagellar mutants or *S. aureus* (**Fig. 4C**).

Biosurfactants properties like charge and toxicity can contribute to their roles as signaling molecules, inhibitors of biofilm formation, and emulsifiers (86). In our experiments, we saw that in contrast to non-ionic Triton X-100 and anionic SDS, addition of the cationic detergent CTAB inhibited motility (**Fig. 3A**) (87), suggesting that charge could impact swarming and surface spreading. Taken together, we posit that biophysical characteristics of surfactants allow surface spreading, but their exact chemical attributes may alter the specifics of the surface motility.

Viscosity, surface hardness, and osmolarity have been previously associated with altering *P. aeruginosa* motility (76, 87). Further, nutrients such as carbon and nitrogen sources as well as iron availability also affect *P. aeruginosa* swarming (27, 35, 39, 88). We observed surface spreading on both semi-solid and hard agar, a condition in which *P. aeruginosa* is nonmotile, however the effect of other physical parameters on this motility are yet to be characterized. While a diverse class of surfactants all transition *P. aeruginosa* to surfing surface motility, it is likely that the combination of physical and chemical cues from the host, neighboring microbial species, and the environment influence this motility and determine the resulting niche expansion and fitness changes in natural habitats.

## MATERIALS AND METHODS

### Bacterial strains and growth conditions

Bacterial strains used in this study are listed in **Supp. Table 1**. *P. aeruginosa* UCBPP-PA14 (89), *S. aureus* JE2 (90), their derivatives, and other species were grown in a modified M63 medium (59) containing 1× M63 salts (13.6 g·L^-1^ KH_2_PO_4_, 2 g·L^-1^ (NH_4_)_2_SO_4_, 0.8 µM ferric citrate, 1 mM MgSO_4_; pH adjusted to 7.0 with KOH) supplemented with 0.3% glucose, 1× ACGU solution (Teknova), 1× supplement EZ (Teknova), 0.1 ng·L^-1^ biotin, and 2 ng·L^-1^ nicotinamide, at 37°C, with shaking at 300 rpm. *V. cholerae* was grown in M63 with 2% NaCl added. *S. aureus* and *P. aeruginosa* clinical isolates from the Cystic Fibrosis Foundation Isolate Core were selected at random from four different patients. *P. aeruginosa* and *S. aureus* transposon mutants were sourced from the PA14NR Set (91) and the Nebraska Transposon Mutant Library (NTML) (90), respectively, and confirmed by PCR using primers flanking the transposon insertion listed in **Supp. Table 7.** For cloning and mutant construction, strains were cultured in Luria Bertani (Miller) broth or on agar plates with 15 g·L^-1^ agar supplemented with 50 µg·mL^-1^ gentamicin and/or 25 µg·mL^-1^ irgasan as needed for selection.

### Preparation of cell-free supernatant

Overnight cultures of bacterial strains were diluted to OD_600_ of 0.05 in fresh media and grown for 24 h in flasks before harvesting at 4000 rpm for 20 minutes. The supernatants were filter-sterilized with a Steriflip with a 0.2 μm polyethersulfone filter (MilliporeSigma). Supernatants were stored at −30°C until use.

### Agar plate motility assays

LB agar plates (for *P. aeruginosa* twitch, and multiple species swim and swarm plates) or M63 agar plates (for *P. aeruginosa* twitch, swim, swarm, and hard agar plates) containing the specified percentage of agar (twitch: 1.5%; swim: 0.3%; swarm: 0.5%; hard: 1.5%) and 25% (v/v) media salts, media salts with the indicated additives, or the indicated species’ supernatant which were mixed into the agar medium, were used for the motility assays. For the twitch assays, M63 plates were used for experiments shown in **Supp. Fig. 1A**, and LB plates for all the other experiments. For the swim assays, LB plates were used for experiments shown in **Supp. Fig. 13**, and M63 plates for all the other experiments. For the swarm assays, LB plates were used for experiments shown in **Supp. Fig. 12 and 14**, and M63 plates for all the other experiments. Freshly poured 25 mL plates were allowed to rest at room temperature for 4 h before inoculation with the specified bacterial species. Twitch plates were prepared prior to the day of the assay. Plates were inoculated as follows: twitch, *P. aeruginosa* colony stabbed through the agar to the bottom of the plate; swim, *P. aeruginosa* or indicated species overnight culture stabbed halfway through the agar; swarm and hard agar, 2 µL of *P. aeruginosa* overnight culture, indicated species, or mixed culture (equal volumes of both strains being mixed) was spotted on the center of the plate. Plates were then incubated for 24 h at 37°C in a single layer of petri dishes in a Mini Low Temperature Incubator (Fisherbrand) for the swarm and surface spreading plates, or for the swim plates, for 24 h or 48 h (as noted) at room temperature, or for the twitch plates, for 48 h at 30°C. When noted, Type II mucin from porcine stomach (Sigma Aldrich), Triton X-100 (Fisher), sodium dodecyl sulfate (SDS) (Fisher BioReagents), hexadecyltrimethylammonium bromide (CTAB) (Sigma Aldrich), surfactin from *B. subtilis* (Sigma Aldrich), rhamnolipids from *P. aeruginosa* (Sigma Aldrich), Mannosylerythritol Lipid A (MEL-A) from *Pseudozyma* yeasts (Cayman), lactonic sophorolipid (Sigma Aldrich), LL-37 (Peptide Sciences), 1,2-Dipalmitoyl-sn-glycero-3-phosphocholine (DPPC) (Sigma Aldrich), and saponin from *Quillaja* bark (Sigma Aldrich) were added at the indicated concentrations to 25% media salts. After incubation, all plates were imaged either using a Chemidoc (Bio-Rad) on the white tray using the Coomassie blue setting or the black tray using the Cy2 (for GFP) or Cy3 (for dsRed or mKO) settings with optimal exposure, or for the twitch plates, using an Epson Perfection V700 Photo scanner with bottom illumination. All experiments were repeated three times.

### Construction of *P. aeruginosa* mutants

Bacterial strains and plasmids are listed in **Supp. Table 1**. For mutant construction, homologous downstream and upstream arms of genes of interest were amplified using the primers listed in **Supp. Table 7**. The amplified fragments were cloned into pDONRPEX18Gm *attP* sites using the Gateway BP Clonase II Enzyme mix (ThermoFisher). Plasmids were transformed into *E. coli* S17-1 λ-pir and confirmed by sequencing prior to conjugation. Conjugants were streaked onto LB plates (without NaCl) + 10% sucrose, and then tested for gentamicin resistance. Gentamicin-sensitive strains were tested for the deletion by PCR and sequencing.

### RNA extraction and library preparation

*P. aeruginosa* was scraped after 17 h from 1.5% or 0.5% agar M63 plates containing 25% (v/v) salts or *S. aureus* cell-free supernatant, resuspended in 2 mL salts mixed with 4 mL of RNAprotect Bacteria Reagent (Qiagen), and then incubated for 5 min at room temperature for stabilization before centrifugation at 4000 rpm for 10 min. Supernatants were completely removed before storage of the pellets at −80°C. RNA was extracted using the Total RNA Purification Plus Kit (Norgen) according to the manufacturer’s instructions for Gram-negative bacteria. The extracted RNA was subjected to an additional genomic DNA removal by DNase I treatment in solution using the TURBO DNA-free Kit (Invitrogen) and checked by PCR for the absence of contaminating DNA. Integrity of the RNA preparation was confirmed by running on an agarose gel and observing intact rRNA bands. Next, rRNA was removed using the *P. aeruginosa* riboPOOLs rRNA Depletion Kit (siTOOLs Biotech) followed by library preparation with the NEBNext Ultra II Directional RNA Library Prep Kit for Illumina (New England Biolabs). The sequencing was performed at the Center for Cancer Research (CCR) Genomics Core Facility. Two biological replicates were collected.

### RNA-seq analysis

The sequencing files were processed with Cutadapt (92) and Trimmomatic (93). Alignment to the *P. aeruginosa* UCBPP-PA14 genome (NCBI) and pairwise comparisons were made using Rockhopper (Wellesley College) (94, 95). Gene expression was set at a minimum baseline value of 4 and *p* values of 0 were changed to the lowest values recorded within a dataset. Ribosomal RNAs and predicted RNAs were removed from each dataset. Upregulated and downregulated genes were based on transcripts that had *p* < 0.05 and log_2_ fold change ≥ 1 or ≤ −1. The location of prophages in the genome of *P. aeruginosa* UCBPP-PA14 NC_008463.1 was determined using PHASTER (96, 97). Venn diagrams were generated using matplotlib_venn package with venn3 using Python (98).

### Gene ontology (GO) enrichment analysis

For *P. aeruginosa* PA14 pathway analysis, the open reading frame designations for the corresponding PAO1 orthologs of *P. aeruginosa* UCBPP-PA14 genes were obtained using the *Pseudomonas* Genome Database (www.pseudomonas.com/rbbh/pairs) (99). The list of designations from the RNA-seq analysis were analyzed at the Gene Ontology Resource (www.geneontology.org) by the PANTHER Overrepresentation Test (released 20221013 and 20231017, as noted) (Annotation Version and Release Date: GO Ontology database DOI: 10.5281/zenodo.7942786 Released 2023-05-10) for enriched biological processes by Fisher’s exact test with Bonferroni correction. All genes identified in the GO enrichment categories ‘heme biosynthesis’ and ‘T3SS’ as well as genes in the same or related neighboring operons, or known to be co-regulated, that were downregulated in both semi-solid and hard agar conditions, are shown in the heatmap in **Fig. 5D**.

#### Accession number(s)

The RNA-seq data have been deposited at NCBI Gene Expression Omnibus (GEO) (https://www.ncbi.nlm.nih.gov/geo/) under accession number GSE249024.

The authors declare no conflict of interest.

## ACKNOWLEDGMENTS

We would like to acknowledge the Center for Cancer Research (CCR) Genomics Core for RNA-sequencing and whole-genome sequencing, the Cystic Fibrosis Foundation Isolate Core for providing clinical isolate strains, and the Adhya, Otto, and Ramamurthi labs at the NIH, the Brinsmade lab at Georgetown University, the Limoli lab at the University of Iowa, and the Goldberg lab at the Emory University School of Medicine for kindly providing bacterial strains. This work utilized the computational resources of the NIH High Performance Computing Biowulf Cluster (http://hpc.nih.gov). The diagrams in Figure 6 was created with Biorender (http://biorender.com). We thank Susan Gottesman, Gisela Storz, Anthony Martini, and Kalinga Pavan Thushara Silva for comments on the manuscript, and members of the Khare, Gottesman, Storz, and Ramamurthi labs for helpful discussion and feedback. This work was supported by the Intramural Research Program of the NIH, National Cancer Institute, Center for Cancer Research. TMZ was supported by a Postdoctoral Research Associate Training (PRAT) Fellowship award 1FI2GM137843-01 from the National Institute of General Medical Sciences.

## COMPETING INTERESTS

No competing interests declared.

## Notes

### Competing Interest Statement

The authors have declared no competing interest.

### Summary of Updates

New data added on motility behavior in other species; manuscript reorganized.

