## Supplemental Figures and Supplemental Tables 1 and 7 for "Interspecies surfactants serve as public goods enabling surface motility in *Pseudomonas aeruginosa*"

**TITLE**

Running head: Surfactants enable motility in *P. aeruginosa*

*Delayna L. Warrell and Tiffany M. Zarrella contributed equally to this work. Author order was determined alphabetically.

Key words: *Pseudomonas aeruginosa*, *Staphylococcus aureus*, surfactants, motility, polymicrobial interactions, phenol-soluble modulins

The authors declare no conflict of interest.

### SUPPLEMENTAL FIGURES AND FIGURE LEGENDS

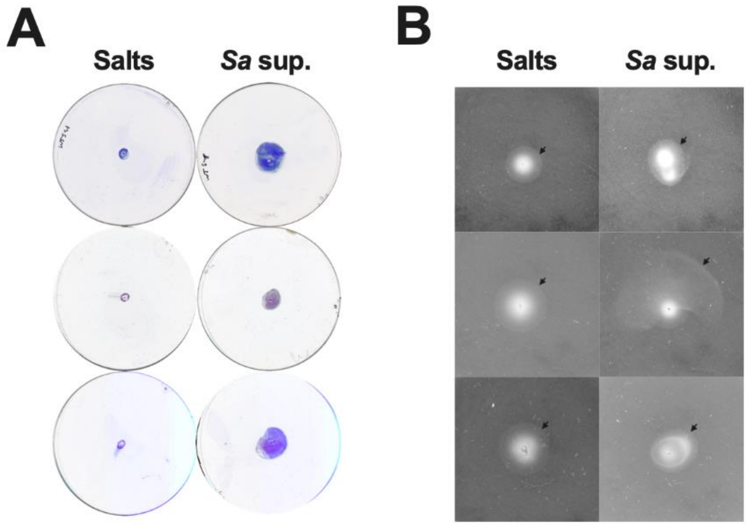

**Supplemental Figure 1. *S. aureus* secreted products alter *P. aeruginosa* twitching and swimming motility.** *P. aeruginosa* was inoculated on **(A)** twitch plates and **(B)** swim plates containing 25% media salts as a control or *S. aureus* supernatant. **(A)** Twitch plates were incubated for 48 hours, visualized by crystal violet staining, and imaged. **(B)** Swim plates were imaged after 24 hours incubation. Arrows indicate the swim boundaries. **(A,B)** Three independent replicates are shown.

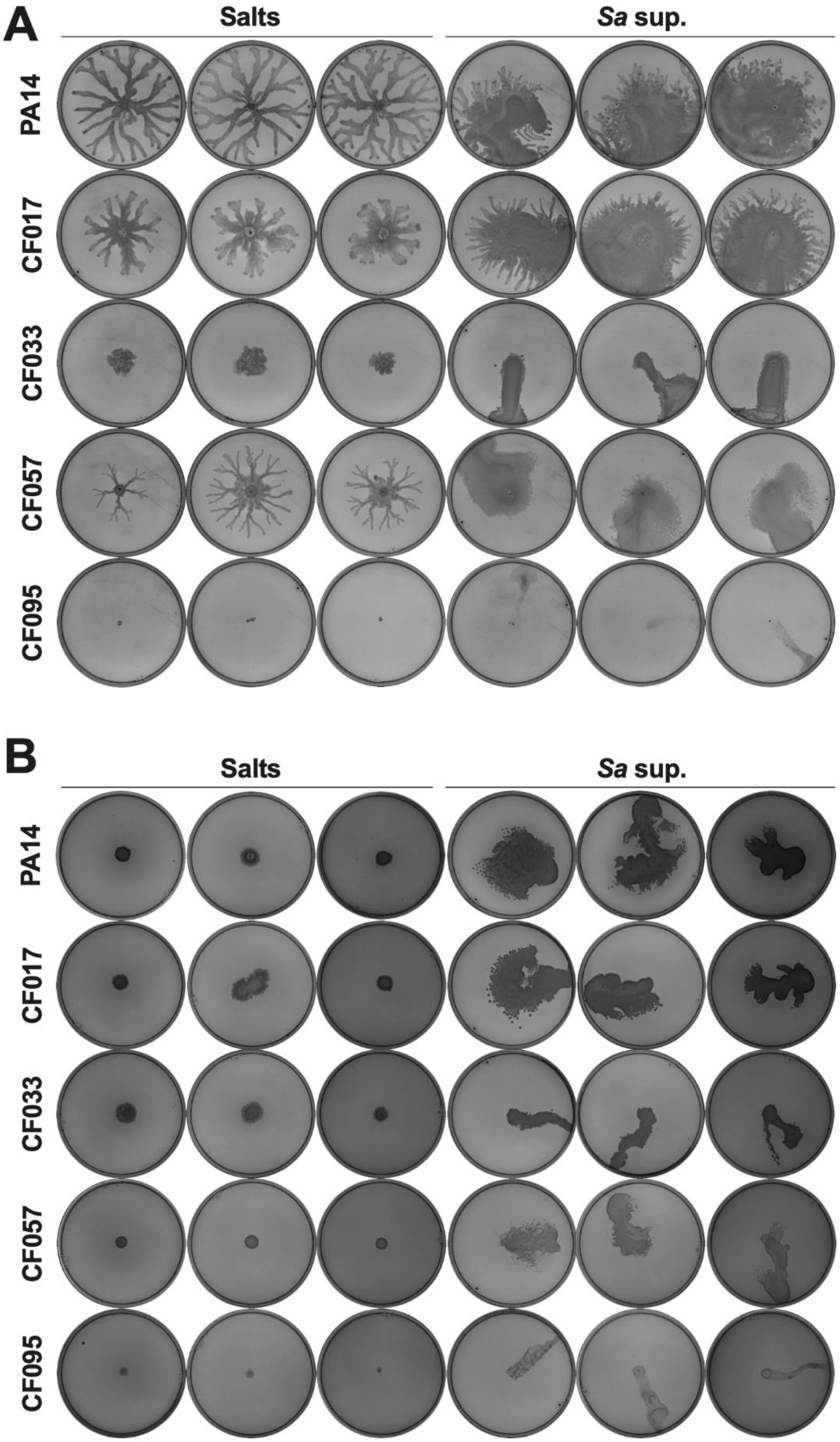
A
Salts
Sa sup.
PA14
CF017
CF033
CF057
CF095
B
Salts
Sa sup.
PA14
CF017
CF033
CF057
CF095

**Supplemental Figure 2. Clinical isolates of *P. aeruginosa* also exhibit surface spreading in the presence of *S. aureus* supernatant.** The indicated *P. aeruginosa* strains were inoculated on **(A)** semi-solid or **(B)** hard agar plates containing 25% media salts as a control or *S. aureus* supernatant, and motility was imaged after 24 hours incubation. **(A,B)** Three independent replicates are shown.

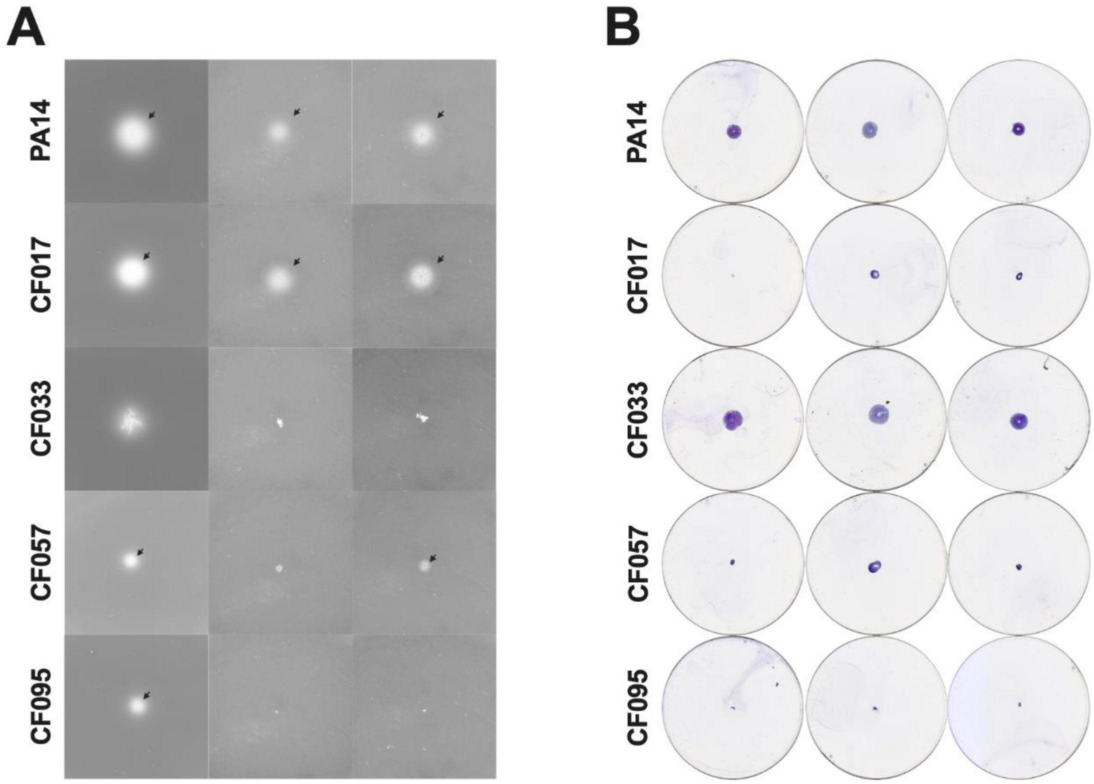

**Supplemental Figure 3. *P. aeruginosa* clinical isolates show motility defects.** The indicated *P. aeruginosa* strains were inoculated on **(A)** swim plates containing 25% media salts as a control or **(B)** LB agar twitch plates. **(A)** Swim plates were imaged after 24 hours incubation. Arrows indicate the swim boundaries. **(B)** Twitch plates were incubated for 48 hours, visualized by crystal violet staining, and imaged. **(A,B)** Three independent replicates are shown.

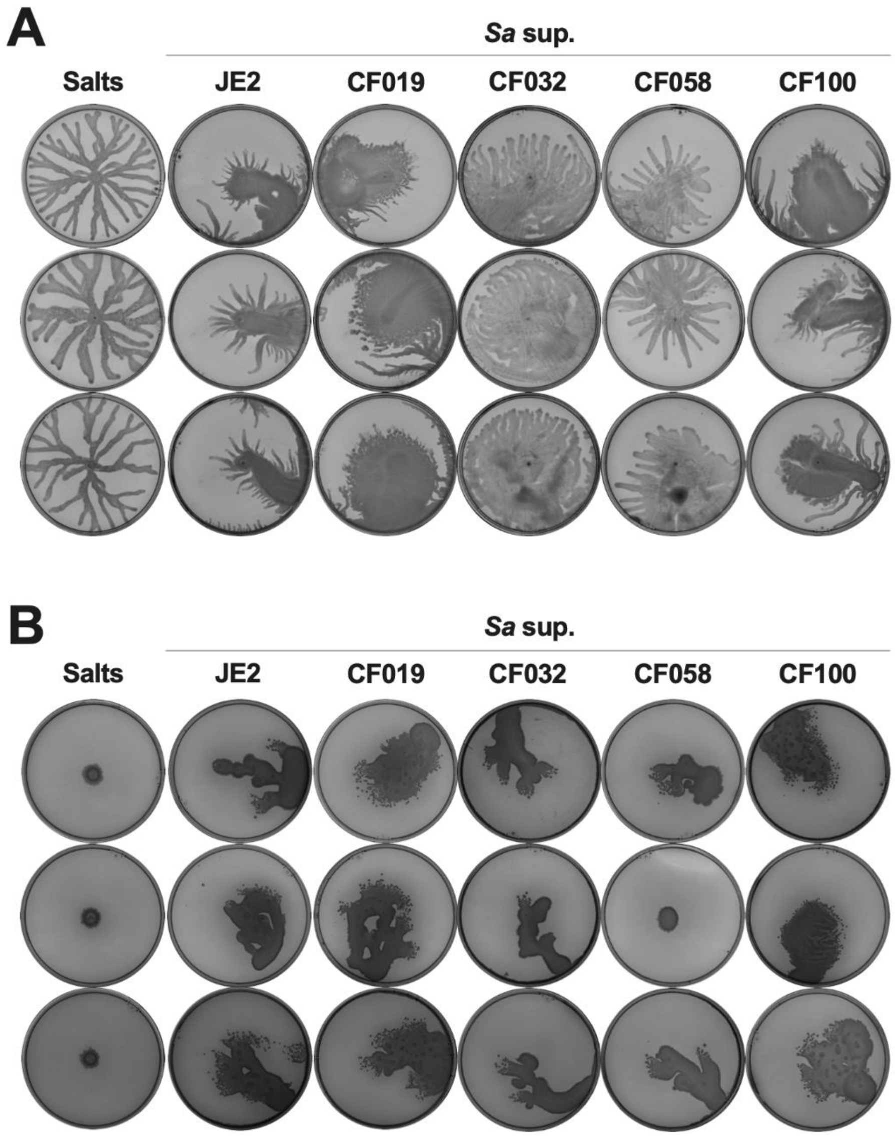

**Supplemental Figure 4. *P. aeruginosa* exhibits surface spreading motility on supernatant from *S. aureus* clinical isolates.** *P. aeruginosa* PA14 was inoculated on **(A)** semi-solid or **(B)** hard agar plates containing 25% media salts as a control or supernatant from the indicated *S. aureus* strains. Motility was imaged after 24 hours incubation. **(A,B)** Three independent replicates are shown.

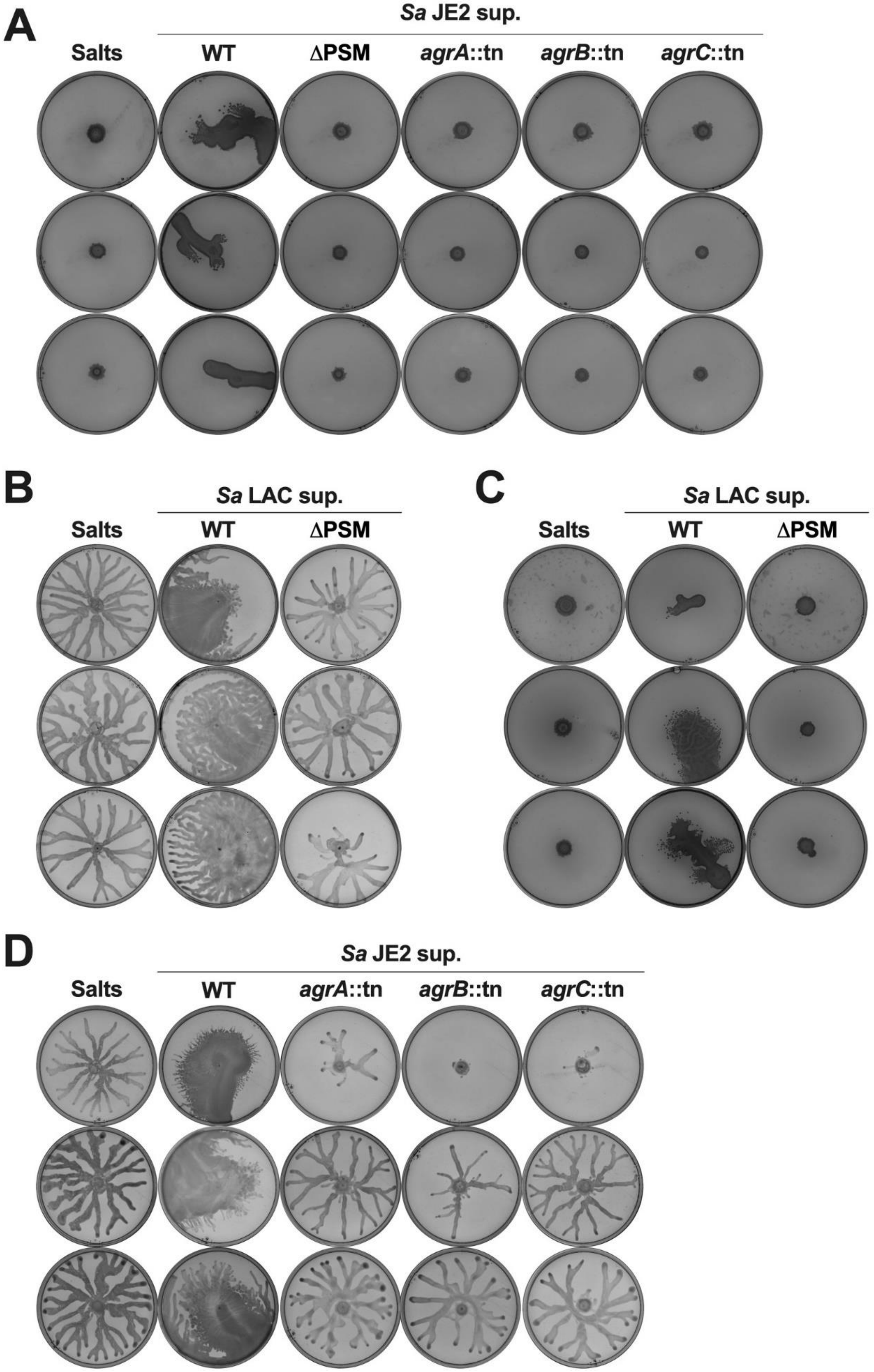

A
Sa JE2 sup.
Salts
WT
ΔPSM
agrA::tn
agrB::tn
agrC::tn
B
Sa LAC sup.
Salts
WT
ΔPSM
C
Sa LAC sup.
Salts
WT
ΔPSM
D
Sa JE2 sup.
Salts
WT
agrA::tn
agrB::tn
agrC::tn

**Supplemental Figure 5. *P. aeruginosa* does not exhibit surface spreading on *S. aureus* supernatant lacking surfactant biosynthesis.** *P. aeruginosa* was inoculated on **(A,C)** hard or **(B,D)** semi-solid agar plates containing 25% media salts as a control or supernatant from the indicated *S. aureus* strains, and motility was imaged after 24 hours incubation. **(A-D)** Three independent replicates are shown.

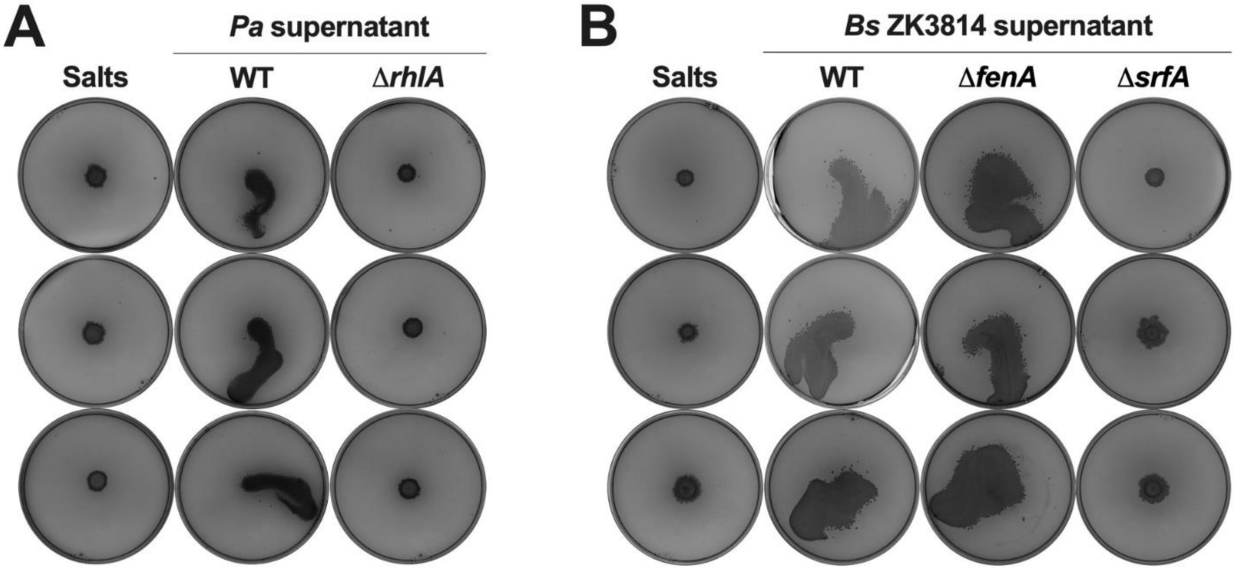

**Supplemental Figure 6. Surfactant biosynthesis is required to facilitate surface spreading in *P. aeruginosa*.** *P. aeruginosa* was inoculated on hard agar plates containing 25% media salts as a control or supernatant from the indicated strain of **(A)** *P. aeruginosa* or **(B)** *B. subtilis* ZK3814, and motility was imaged after 24 hours incubation. **(A,B)** Three independent replicates are shown.

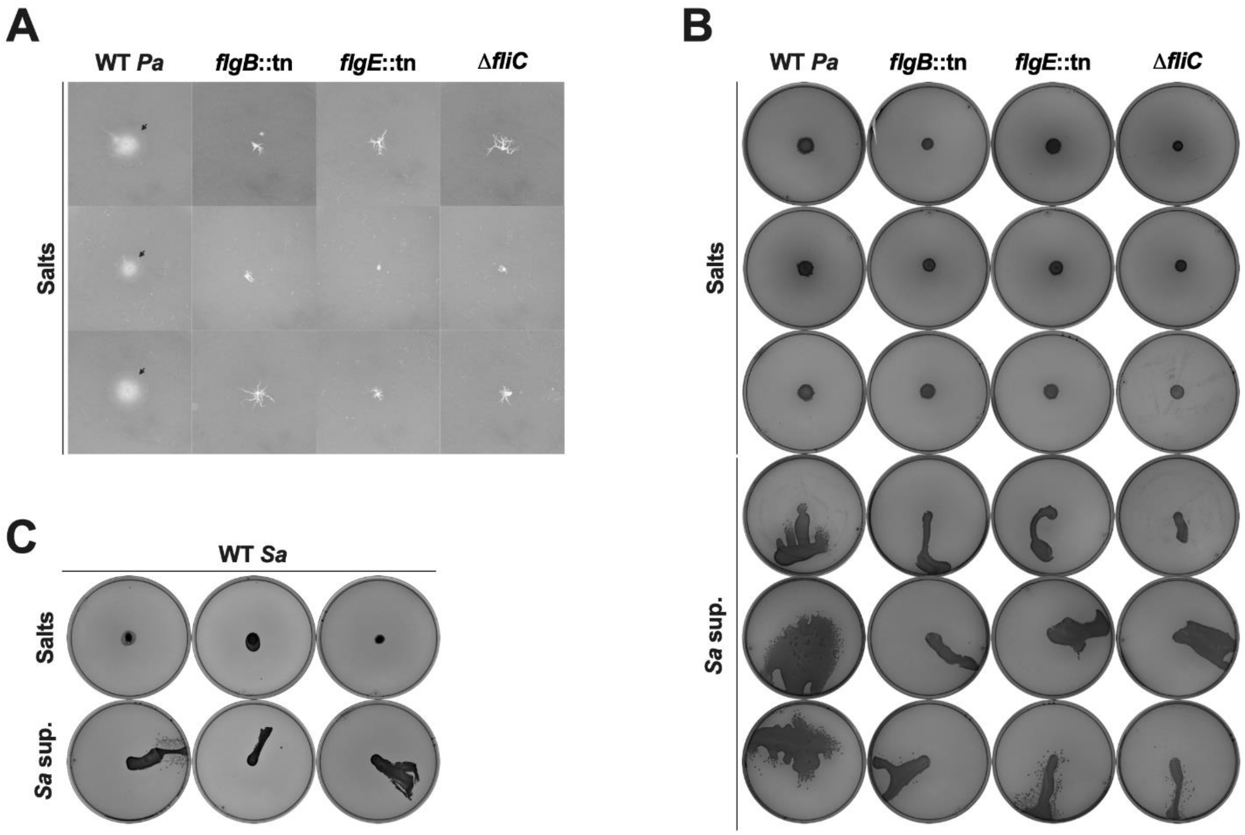

**Supplemental Figure 7. *P. aeruginosa* flagellar mutants do not exhibit surface spreading motility.** The indicated strains of **(A,B)** *P. aeruginosa* or **(C)** *S. aureus* were inoculated on **(A)** swim plates or **(B,C)** hard agar plates containing 25% media salts as a control or *S. aureus* supernatant, and motility was imaged after 24 hours incubation. **(A)** Arrows indicate the swim boundaries. **(A-C)** Three independent replicates are shown.

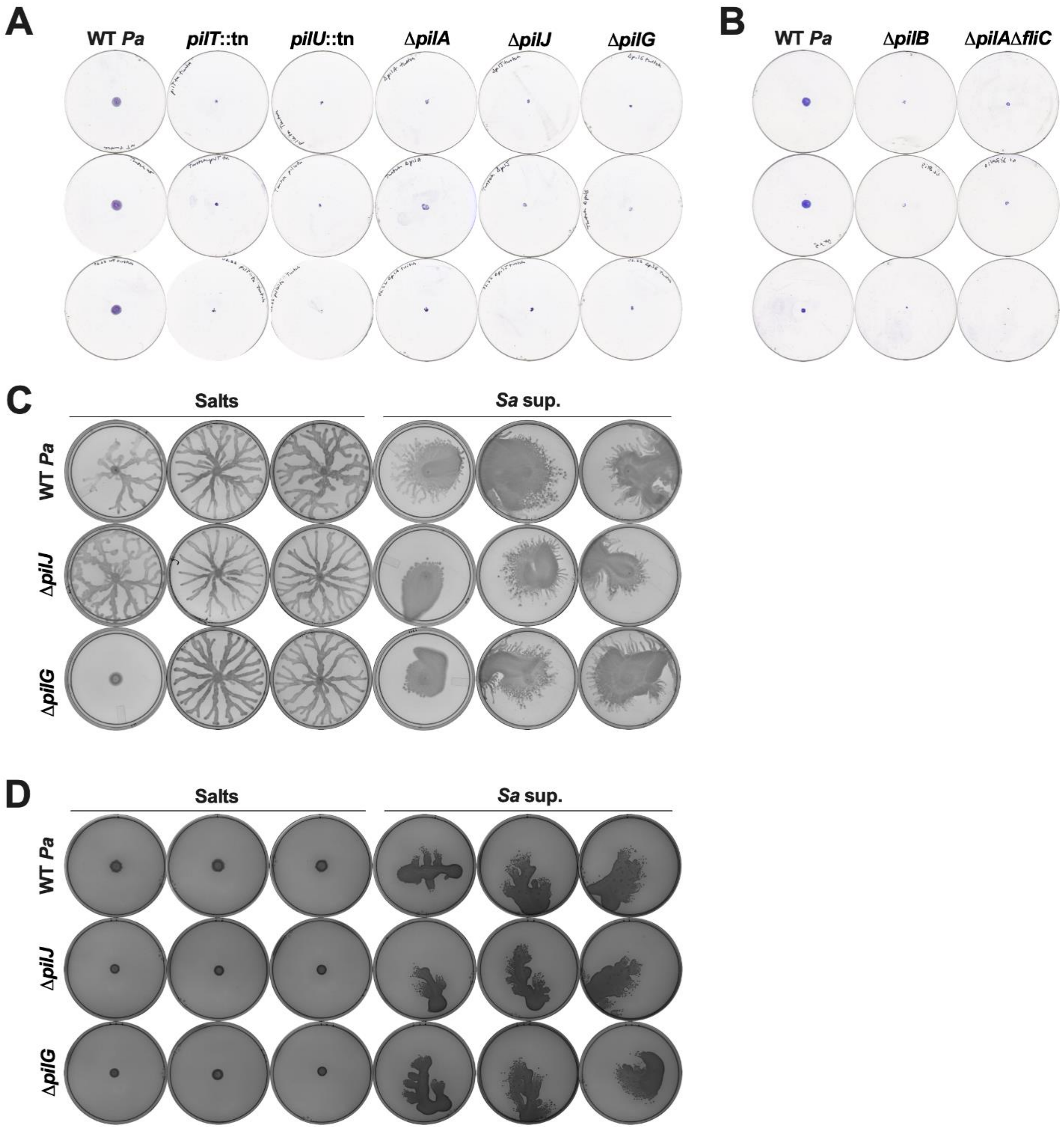

**Supplemental Figure 8. *P. aeruginosa* pili mutants do not twitch but exhibit surface spreading. (A)** Twitch plates composed of LB agar were inoculated with the indicated strain of *P. aeruginosa*, incubated for 48 hours, visualized by crystal violet staining, and imaged. **(B,C)** The indicated *P. aeruginosa* strains were inoculated on **(B)** semi-solid or **(C)** hard agar plates containing 25% media salts as a control or *S. aureus* supernatant as indicated. **(B,C)** Plates were imaged after 24 hours incubation. **(A-C)** Three independent replicates are shown.

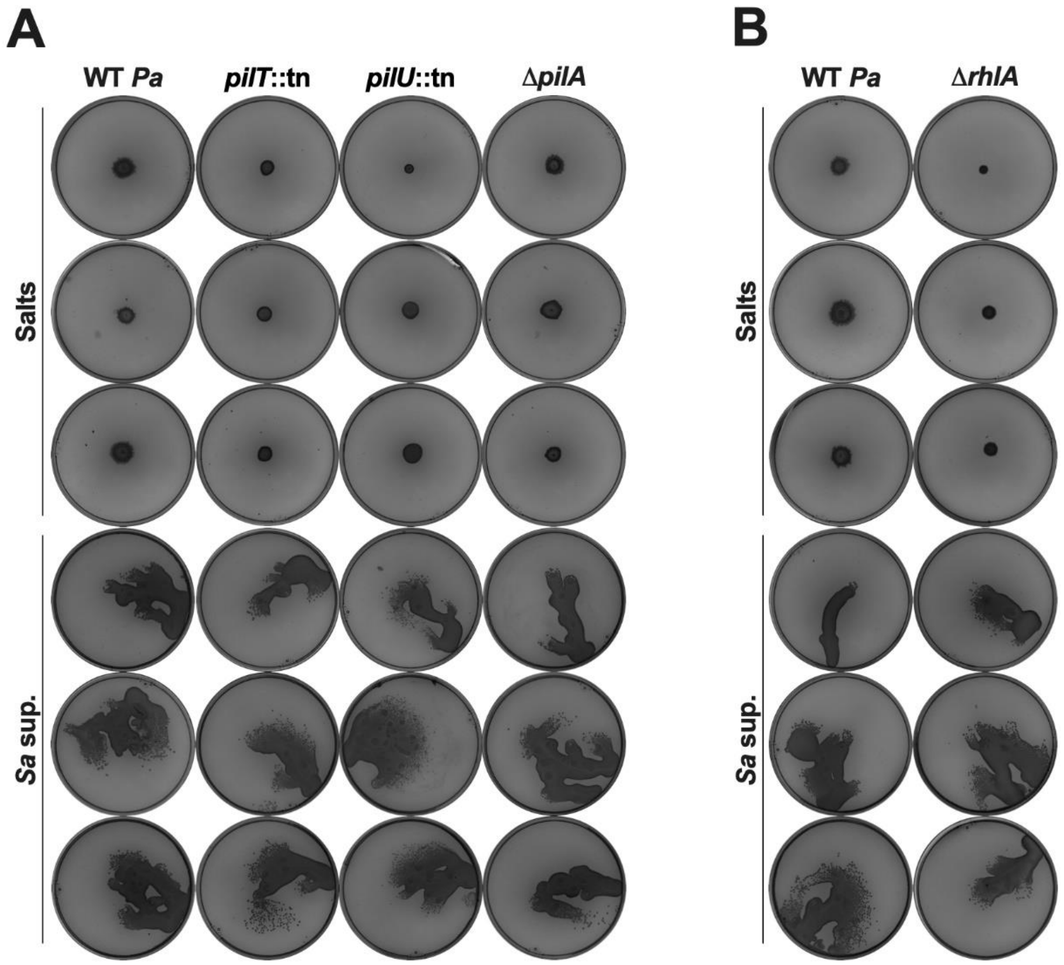

**Supplemental Figure 9. Pili function and rhamnolipids are not required for surface spreading on hard agar. (A,B)** The indicated *P. aeruginosa* strains were inoculated on hard agar plates containing 25% media salts as a control or *S. aureus* supernatant, and motility was imaged after 24 hours incubation. Three independent replicates are shown.

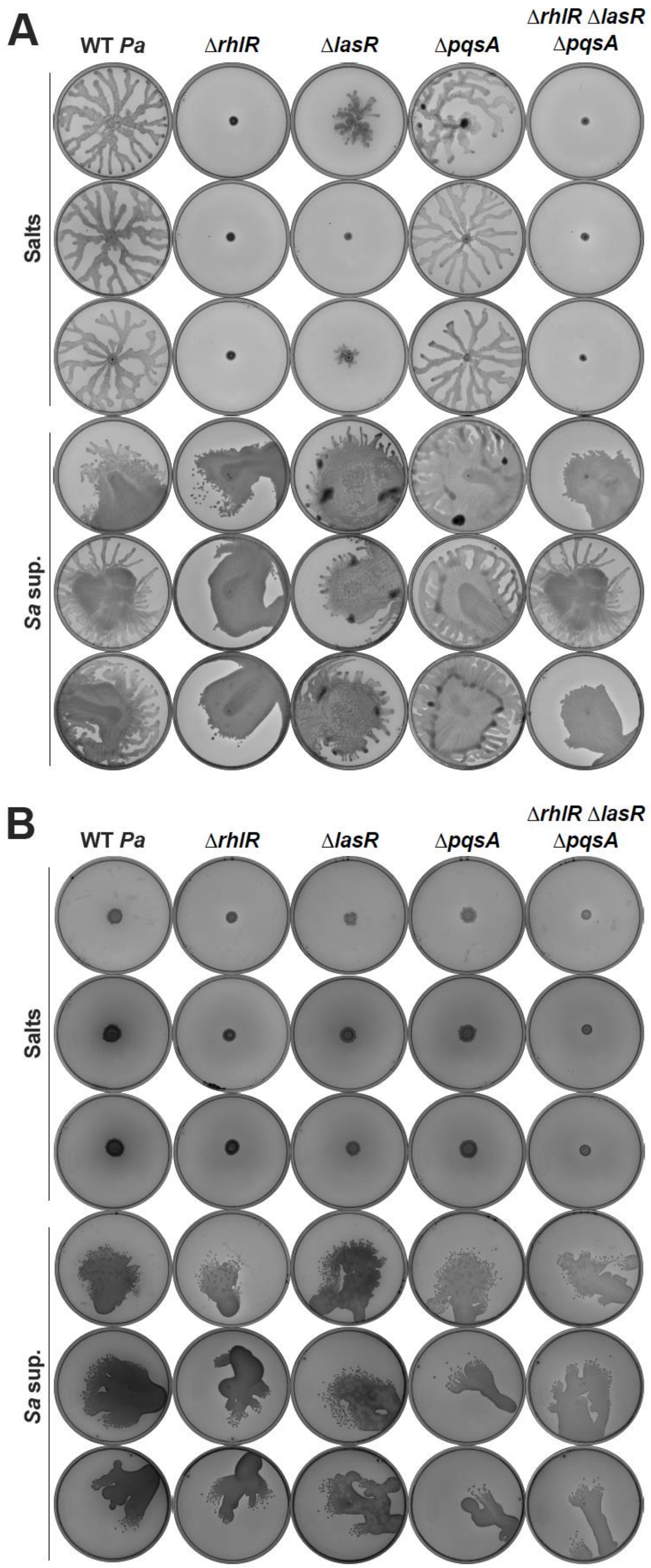
A
WT Pa
ΔrhlR
ΔlasR
ΔpqsA
ΔrhlR ΔlasR ΔpqsA
Salts
Sa sup.
B
WT Pa
ΔrhlR
ΔlasR
ΔpqsA
ΔrhlR ΔlasR ΔpqsA
Salts
Sa sup.

**Supplemental Figure 10. The *P. aeruginosa* quorum sensing systems are not required for surface spreading motility.** The indicated *P. aeruginosa* strains were inoculated on **(A)** semi-solid or **(B)** hard agar plates containing 25% media salts as a control or *S. aureus* supernatant, and motility was imaged after 24 hours incubation. **(A,B)** Three independent replicates are shown.

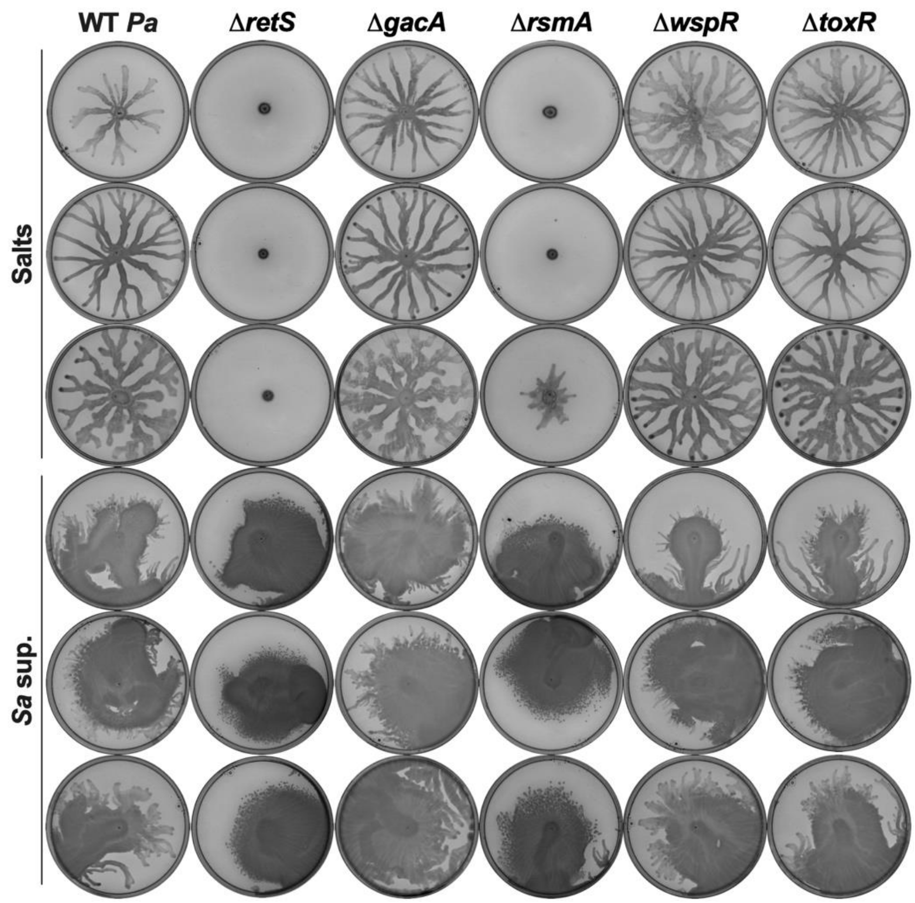

**Supplemental Figure 11. Several *P. aeruginosa* regulators are not required for surface spreading motility.** The indicated *P. aeruginosa* strains were inoculated on semi-solid agar plates containing 25% media salts as a control or *S. aureus* supernatant, and motility was imaged after 24 hours incubation. Three independent replicates are shown.

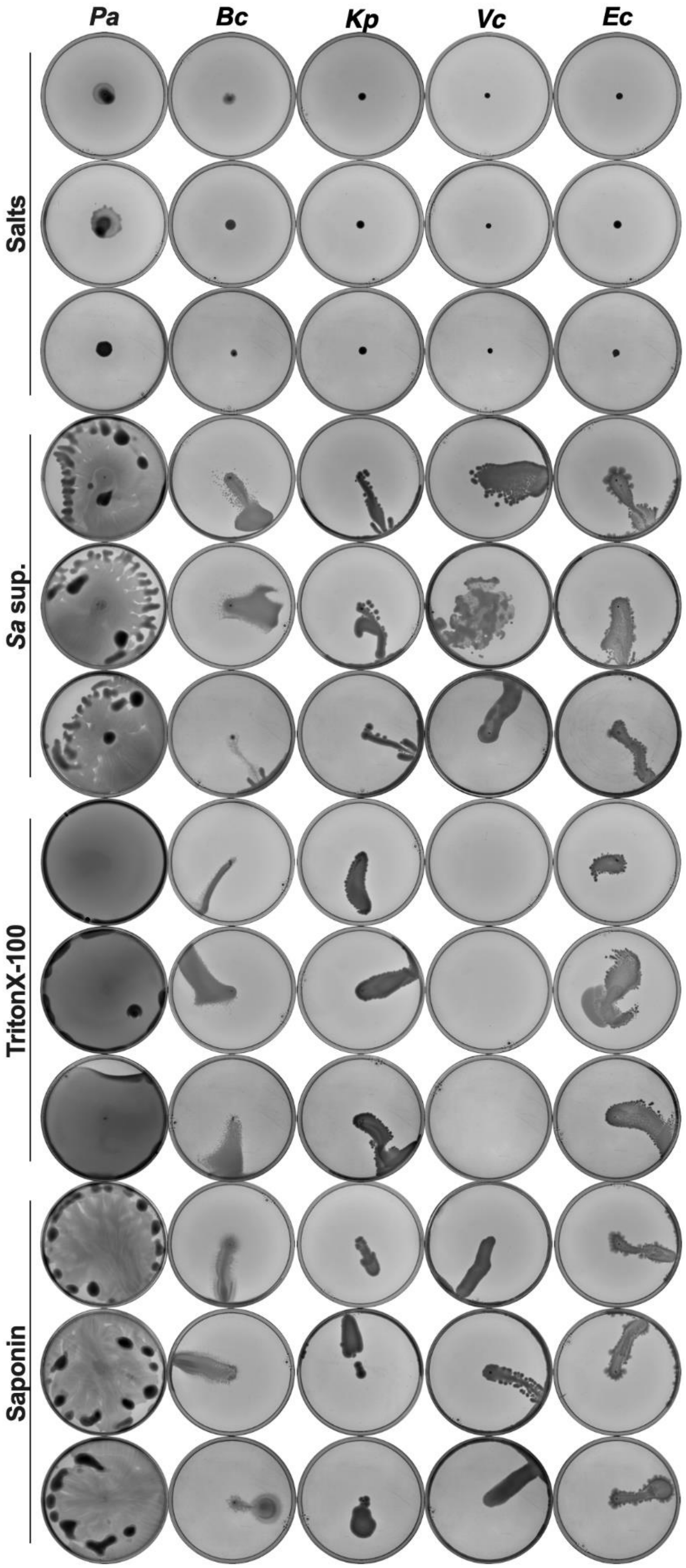
Pa
Bc
Kp
Vc
Ec
Salts
Sa sup.
TritonX-100
Saponin

**Supplemental Figure 12. Surface spreading is not a universal bacterial response to exogenous surfactants.** The indicated species – *P. aeruginosa* PA14, *Burkholderia cenocepacia* K56-2, *K. pneumoniae*, *V. cholerae*, or *E. coli* – were inoculated on semi-solid LB agar plates containing 25% media salts as a control, *S. aureus* supernatant, or 25% media salts with the addition of 0.1% Triton X-100 or 25 $\mu g \cdot mL^{-1}$ saponin and motility was imaged after 24 hours incubation. Three independent replicates are shown.

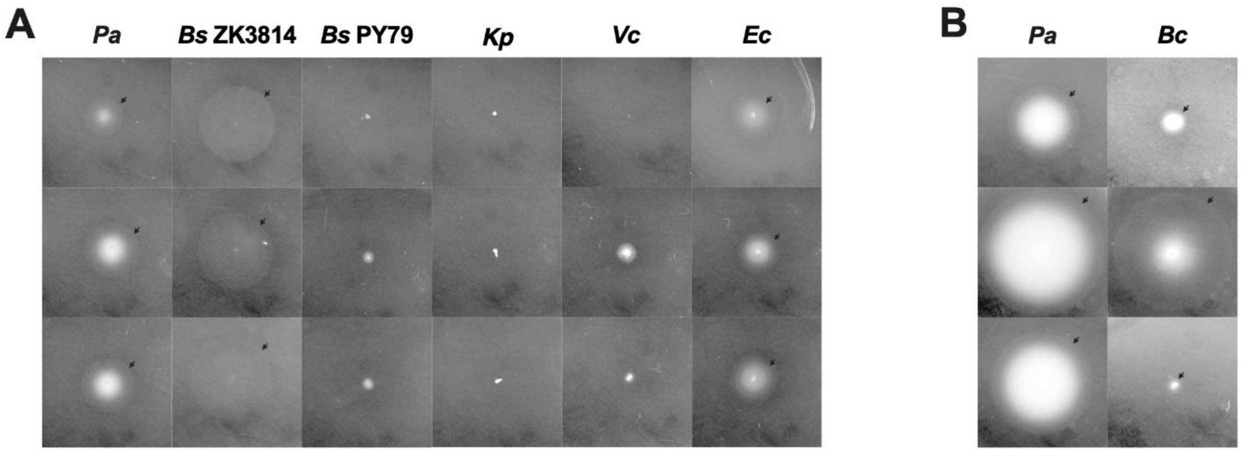

**Supplemental Figure 13. *P. aeruginosa*, *B. subtilis* ZK3814, *E. coli*, and *B. cenocepacia* demonstrate swimming motility. (A,B)** The indicated species – *P. aeruginosa* PA14, *B. subtilis* ZK3814 and PY79, *B. cenocepacia* K56-2, *K. pneumoniae*, *V. cholerae*, or *E. coli* – were inoculated on LB swim plates containing 25% media salts as a control. Plates were imaged after **(A)** 24 hours or **(B)** 48 hours incubation. **(A,B)** Arrows indicate the swim boundaries. Three independent replicates are shown.

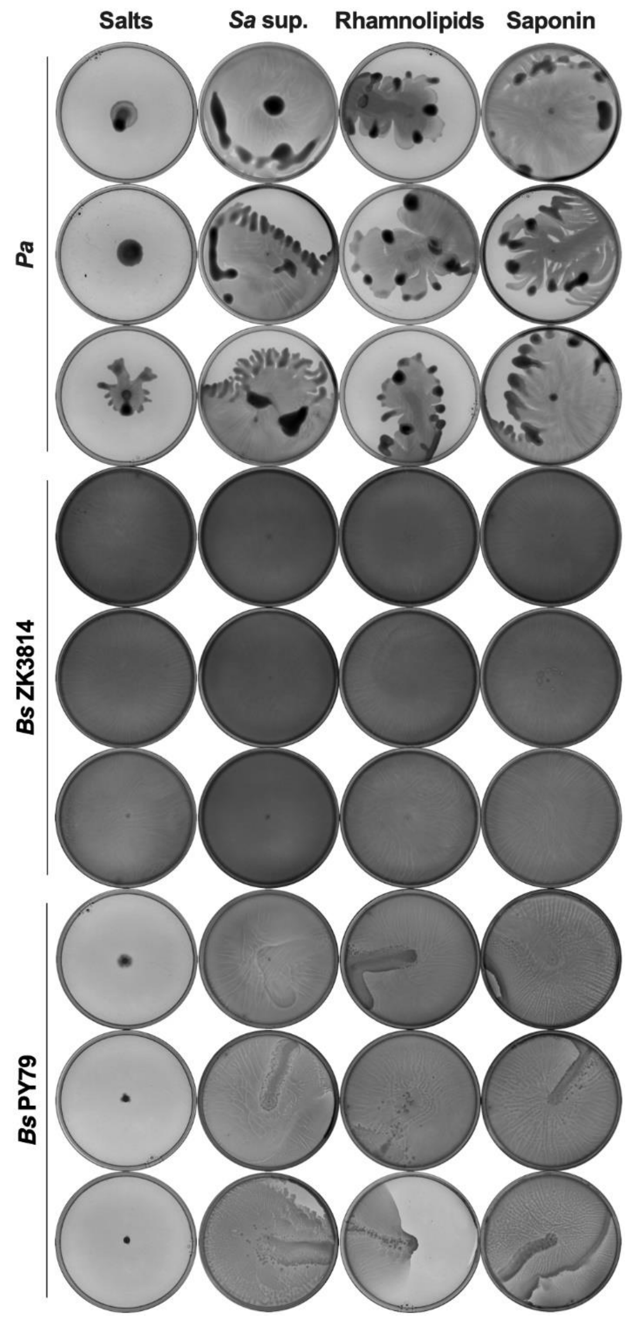

**Supplemental Figure 14. Exogenous surfactants restore swarming in surfactant-deficient *B. subtilis*.** The indicated species – *P. aeruginosa* PA14 or *B. subtilis* ZK3814 and PY79 – were inoculated on semi-solid LB agar plates containing 25% media salts as a control, *S. aureus* supernatant, or 25% media salts with the addition of 50 $\mu g \cdot mL^{-1}$ rhamnolipids or 25 $\mu g \cdot mL^{-1}$ saponin and motility was imaged after 24 hours incubation. Three independent replicates are shown.

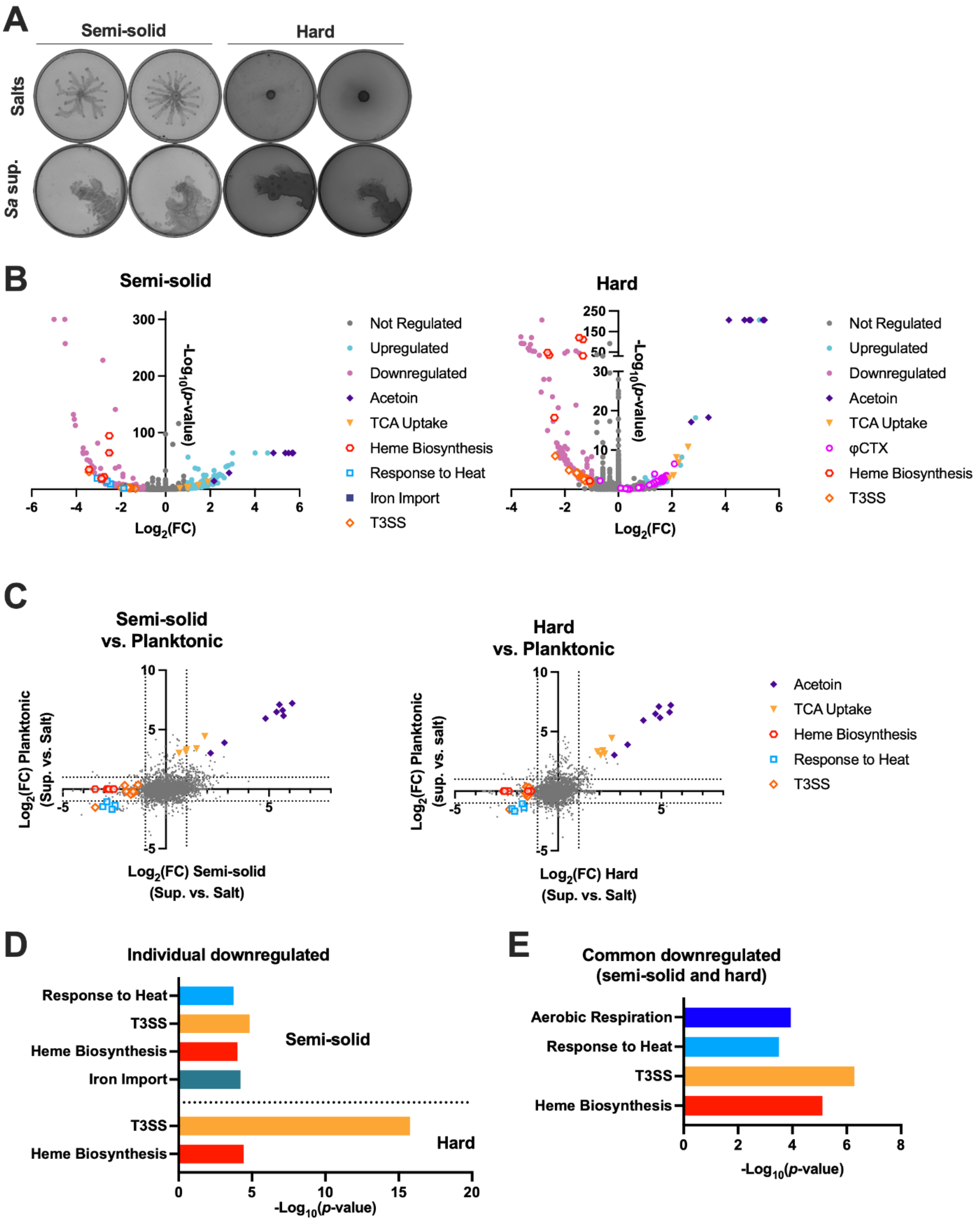

**Supplemental Figure 15.** ***P. aeruginosa*** **differentially regulates genes when undergoing surface spreading vs. planktonic growth in the presence of** ***S. aureus*** **secreted products.**

**(A)** WT *P. aeruginosa* cells were scraped from the leading edge on semi-solid and hard agar plates containing 25% media salts as a control or *S. aureus* supernatant after 17 hours of incubation. Both independent replicates used for the RNA-seq are shown prior to scraping. **(B)** Volcano plots of -$\log_{10}$($p$-value) vs. $\log_2$(fold-change) transcript levels after exposure to *S. aureus* supernatant compared to media salts control after 17 hours on **(left)** semi-solid or **(right)** hard agar. Genes shown as upregulated or downregulated have $p < 0.05$ and $\log_2$(fold-change) $\geq 1$ or $\leq -1$ respectively. **(C)** Scatter plots of $\log_2$(fold-change) transcript levels in *P. aeruginosa* after 17 hours on **(left)** semi-solid or **(right)** hard agar containing *S. aureus* supernatant compared to 2 hours exposure to *S. aureus* supernatant in planktonic culture (1). Genes shown as upregulated or downregulated have $p < 0.05$ and $\log_2$(fold-change) $\geq 1$ or $\leq -1$ respectively. **(D,E)** Gene Ontology (GO) enrichment (2-4) of *P. aeruginosa* genes downregulated after 17 hours incubation on **(D, top)** semi-solid and **(D, bottom)** hard agar plates or commonly downregulated on **(E)** both agar plates containing *S. aureus* supernatant. Nonredundant categories shown.

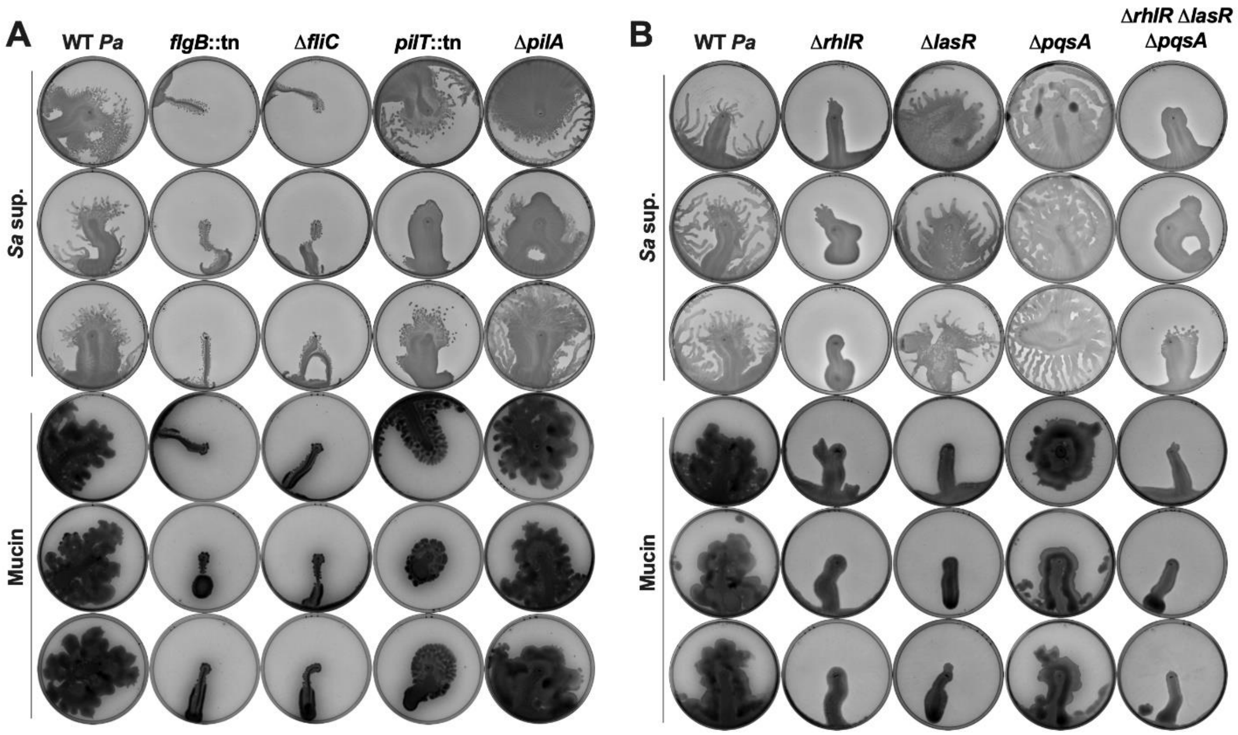

**Supplemental Figure 16. Quorum sensing and flagella, but not pili, are required for motility on mucin. (A,B)** The indicated *P. aeruginosa* strains were inoculated on semi-solid agar plates containing 25% *S. aureus* supernatant or 25% media salts with the addition of 0.4% mucin, and motility was imaged after 24 hours incubation. Three independent replicates are shown.

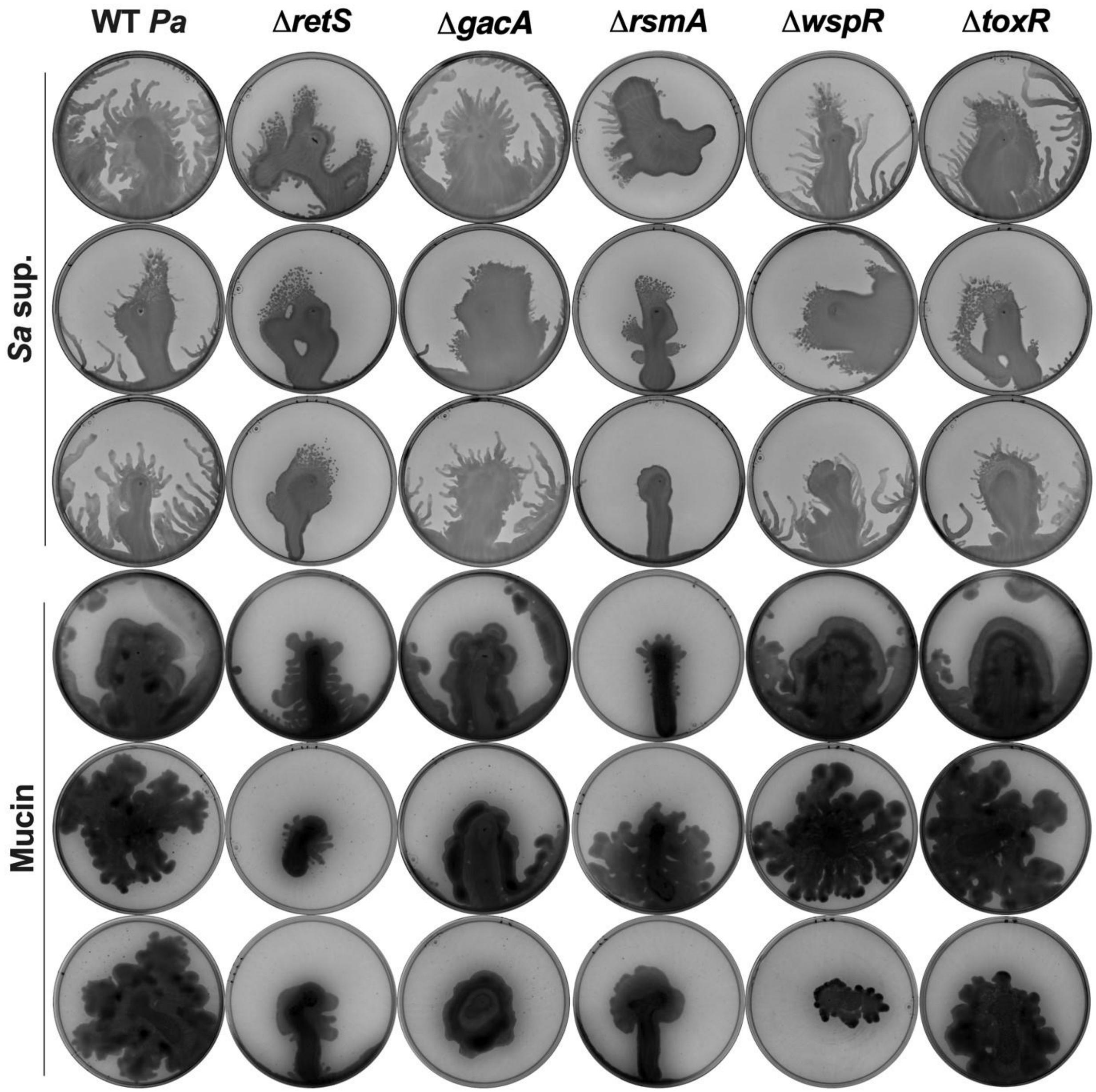

**Supplemental Figure 17. Known *P. aeruginosa* motility regulatory systems are not required for movement on mucin.** The indicated *P. aeruginosa* strains were inoculated on semi-solid agar plates containing 25% of *S. aureus* supernatant or 25% media salts with the addition of 0.4% mucin, and images were taken after 24 hours incubation. Three independent replicates shown.

### SUPPLEMENTAL TABLES

**Supp. Table 1. Bacterial strains and plasmids used in this study.**

| Strain | Description[a,b] | Source |
|---|---|---|
| *P. aeruginosa* | | |
| PA14 | University of California Berkeley Plant Pathology (UCBPP)-PA14 | (5) |
| 33660 | PA14 MAR2xT7 PA14_05180::tn (*pilT*); Gent$^r$ | (6) |
| 42562 | PA14 MAR2xT7 PA14_50480::tn (*flgB*); Gent$^r$ | (6) |
| 48241 | PA14 MAR2xT7 PA14_50450::tn (*flgE*); Gent$^r$ | (6) |
| 53607 | PA14 MAR2xT7 PA14_05190::tn (*pilU*); Gent$^r$ | (6) |
| AK619 | PA14 ΔPA14_51430 (*pqsA*) unmarked | (7) |
| CF017 | AMT0194-10; co-isolated with CF019; age: 4.91 years | CFF Isolate Core |
| CF033 | AMT0457-07; co-isolated with CF032; age: 4.28 years | CFF Isolate Core |
| CF057 | AMT0461-12; co-isolated with CF058; age: 12.16 years | CFF Isolate Core |
| CF095 | AMT0492-16; co-isolated with CF100; age: 13.52 years | CFF Isolate Core |
| SB049 | PA14 expressing GFP; Gent$^r$ | This study |
| SB092 | PA14 ΔPA14_19100 (*rhlA*) unmarked | This study |
| SB387 | PA14 ΔPA14_45960 (*lasR*) unmarked | This study |
| SB392 | PA14 ΔPA14_19120 (*rhlR*) unmarked | This study |
| SB400 | PA14 ΔPA14_05360 (*pilJ*) unmarked | This study |
| SB401 | PA14 ΔPA14_05320 (*pilG*) unmarked | This study |
| SB402 | PA14 ΔPA14_16500 (*wspR*) unmarked | This study |
| SB403 | PA14 ΔPA14_55160 (*toxR*) unmarked | This study |
| SB426 | PA14 ΔPA14_30650 (*gacA*) unmarked | This study |
| SB428 | PA14 ΔPA14_58730 (*pilA*) unmarked | This study |
| SB431 | PA14 ΔPA14_64230 (*retS*) unmarked | This study |
| SB446 | PA14 ΔPA14_50290 (*fliC*) unmarked | This study |
| SB510 | PA14 ΔPA14_58750 (*pilB*) unmarked | This study |
| SB518 | PA14 ΔPA14_50290 (*fliC*) ΔPA14_58730 (*pilA*) unmarked | This study |
| SB452 | PA14 ΔPA14_45960 (*lasR*) ΔPA14_19120 (*rhlR*) ΔPA14_51430 (*pqsA*) unmarked | This study |
| SB454 | PA14 ΔPA14_52570 (*rsmA*) unmarked | This study |
| SB485 | SB446 expressing GFP | This study |
| SB489 | PA14 expressing mKO | This study |
| *S. aureus* | | |
| JE2 | *Staphylococcus aureus* subsp. *aureus* USA300_FPR3757 (CA-MRSA)-JE2 | (8) |
| NE95 | JE2 SAUSA300_1989::tn (*agrB*); Em$^r$ | (9) |
| NE873 | JE2 SAUSA300_1991::tn (*agrC*); Em$^r$ | (9) |
| NE1532 | JE2 SAUSA300_1992::tn (*agrA*); Em$^r$ | (9) |
| CF019 | AMT0194-12; co-isolated with CF017; age: 4.91 years | CFF Isolate Core |
| CF032 | AMT0457-03; co-isolated with CF033; age: 4.28 years | CFF Isolate Core |
| CF058 | AMT0461-13; co-isolated with CF057; age: 12.16 years | CFF Isolate Core |
| CF100 | AMT0492-21; co-isolated with CF095; age: 13.52 years | CFF Isolate Core |
| SB209 | JE2 pKM16; Cm$^r$ | This study |

| | | |
|---|---|---|
| SB350 | JE2 ΔPSM alpha1-4 delta (ATT) | D. Limoli; (10) |
| SB371 | LAC | M. Otto |
| SB372 | LAC ΔPSM alpha1-4 beta1-2 delta (ATT) | M. Otto; (11) |
| *E. coli* | | |
| *ccdB* Survival 2 T1$^R$ | *E. coli* strain used for maintenance of pDONR plasmid | Invitrogen |
| DH5α | *E. coli* strain used for cloning | NEB |
| IM08B | *E. coli* strain used for cloning | BEI Resources |
| S17-1 λ-pir | *E. coli* strain used for conjugation | (12) |
| AK111 | MG1655 SB144 | (13) |
| Other species | | |
| SB145 | *Bacillus subtilis* PY79 | K. Ramamurthi |
| SB146 | *Burkholderia cenocepacia*; ATCC 25608 | S. Adhya |
| SB147 | *Klebsiella pneumoniae* subsp. *pneumoniae* KPNIH1 | S. Adhya |
| SB149 | *Vibrio cholerae* | S. Adhya |
| SB373 | *B. subtilis* ZK3814 | M. Otto; (14) |
| SB374 | ZK3814 Δ*fenA* | M. Otto; (14) |
| SB375 | ZK3814 Δ*srfA* | M. Otto; (14) |
| SB468 | *B. cenocepacia*; ATCC BAA-245 | S. Adhya |
| SB480 | *B. cenocepacia* K56-2 | J. Goldberg |
| **Plasmids** | | |
| *Gene deletion* | | |
| pDONRPEX18Gm | Shuttle vector with *attP* sites and *ccdB*; Cm$^r$ Gent$^r$ | (15) |
| pAK611 | pEX18ApGW: ΔPA14_51430 (*pqsA*); Ap$^r$ Gent$^r$ | (7) |
| pSB445 | pDONRPEX18Gm: ΔPA14_50290 (*fliC*); Gent$^r$ | This study |
| pSB491 | pDONRPEX18Gm: ΔPA14_19100 (*rhlA*); Gent$^r$ | This study |
| pSB388 | pDONRPEX18Gm: ΔPA14_45960 (*lasR*); Gent$^r$ | This study |
| pSB391 | pDONRPEX18Gm: ΔPA14_19120 (*rhlR*); Gent$^r$ | This study |
| pSB396 | pDONRPEX18Gm: ΔPA14_05360 (*pilJ*); Gent$^r$ | This study |
| pSB397 | pDONRPEX18Gm: ΔPA14_05320 (*pilG*); Gent$^r$ | This study |
| pSB398 | pDONRPEX18Gm: ΔPA14_16500 (*wspR*); Gent$^r$ | This study |
| pSB399 | pDONRPEX18Gm: ΔPA14_55160 (*toxR*); Gent$^r$ | This study |
| pSB420 | pDONRPEX18Gm: ΔPA14_64230 (*retS*); Gent$^r$ | This study |
| pSB421 | pDONRPEX18Gm: ΔPA14_30650 (*gacA*); Gent$^r$ | This study |
| pSB422 | pDONRPEX18Gm: ΔPA14_58730 (*pilA*); Gent$^r$ | This study |
| pSB453 | pDONRPEX18Gm: ΔPA14_52570 (*rsmA*); Gent$^r$ | This study |
| pSB509 | pDONRPEX18Gm:ΔPA14_58750 (*pilB*); Gent$^r$ | This study |
| *Express fluorescent marker* | | |
| pKM16 | Expressing DsRed from SarA-P1 promoter; Ap$^r$ Cm$^r$ | S. Brinsmade; (16) |
| pUC18T-mini-Tn7T- | Expressing GFP; Ap$^r$ Gent$^r$ | H. Schweizer; (17) |

| | | |
|---|---|---|
| Gm-*gfpmut3* | | |
| pUC18T-mini-Tn7T-Gm-*mKO* | Expressing mKO; Ap$^r$ Gent$^r$ | H. Schweizer; (17) |

[a] Ap$^r$, ampicillin resistance (*E. coli*); Cm$^r$, chloramphenicol resistance (*E. coli*, *S. aureus*); Em$^r$, erythromycin resistance (*S. aureus*); Gent$^r$, gentamicin resistance (*P. aeruginosa*)

[b] CFF Isolate Core Samples are annotated as follows: AMT####-## (Patient ID-Isolate number)

**Supp. Table 2. Gene expression in *P. aeruginosa* from semi-solid or hard agar at 17 hours after exposure to *S. aureus* supernatant or media salts control determined from RNA-seq analysis. (Excel)** Gene expression was analyzed by RNA-seq from RNA purified from two biological replicates of each treatment at 17 hours (semi-solid or hard agar with addition of *S. aureus* supernatant or media salts control).

**Supp. Table 3. Differentially expressed genes between *P. aeruginosa* from semi-solid or hard agar plates after exposure to *S. aureus* supernatant or media salts control determined from RNA-seq analysis. (Excel)** Gene expression was analyzed by RNA-seq from purified RNA from two biological replicates of each treatment after 17 hours (semi-solid or hard agar with addition of *S. aureus* supernatant or media salts control). Fold change indicates the mean expression of the supernatant-exposed *P. aeruginosa* over the control.

**Supp. Table 4. Differentially expressed genes in common between *P. aeruginosa* from semi-solid and hard agar plates after exposure to *S. aureus* supernatant or media salts control determined from RNA-seq analysis. (Excel)** Gene expression was analyzed by RNA-seq from purified RNA from two biological replicates of each treatment after 17 hours (semi-solid or hard agar with addition of *S. aureus* supernatant or media salts control). Genes commonly upregulated or downregulated from the semi-solid and hard agar conditions are listed. Fold change indicates the mean expression of the supernatant-exposed *P. aeruginosa* over the control.

**Supp. Table 5. Differentially expressed genes in common between *P. aeruginosa* from semi-solid or hard agar and planktonic growth, or semi-solid and hard agar exclusively, after exposure to *S. aureus* supernatant or media salts control determined from RNA-seq analysis. (Excel)** Gene expression was analyzed by RNA-seq from purified RNA from two biological replicates of each treatment after 17 hours (semi-solid or hard agar and planktonic growth with addition of *S. aureus* supernatant or media salts control). Genes commonly upregulated or downregulated from the semi-solid or hard agar and planktonic growth conditions, or semi-solid and hard agar exclusively, are listed. Fold change indicates the mean expression of the supernatant-exposed *P. aeruginosa* over the control.

**Supp. Table 6. Gene ontology (GO) enrichment of upregulated and downregulated genes after *P. aeruginosa* exposure to *S. aureus* exoproducts. (Excel)** GO enrichment of *P. aeruginosa* genes differentially expressed on semi-solid and hard agar plates individually as well as in both, in common with planktonic cells, as well as exclusively in both motility conditions (but not in planktonic cells).

**Supp. Table 7. Primers used in this study*.**

| Primer | Sequence 5′→3′ | Site^ | Location* | Application |
|---|---|---|---|---|
| Pa003 | ggacttatcagccaacctgtt | - | MAR2xT7 transposon | Verify transposon insertion |
| Pa061 | ggggacaagtttgtacaaaaaagcaggctcaccatgtgaccctcgagttct | *attB1* | Up PA14_19100 | Generate mutant |
| Pa062 | ggttcaggcgtagccgatcatctcacacctcccaaaaa | - | Up and down PA14_19100 | Generate mutant |
| Pa063 | tttttgggaggtgtgagatgatcggctacgcctgaacc | - | Up and down PA14_19100 | Generate mutant |
| Pa064 | ggggaccactttgtacaagaaagctgggtacacgttgaacttggggtgta | *attB2* | Down PA14_19100 | Generate mutant |
| Pa065[#] | aaatcggacaagtggattcg | - | Up PA14_19100 | Sanger sequencing |
| Pa066[#] | atcgagaaagcgttgcagtt | - | Down PA14_19100 | Sanger sequencing |
| Pa108 | tgaagaaccaggacccgatg | | Up PA14_50450 | Verify transposon insertion |
| Pa109 | gctcatggagacgtacagca | | Down PA14_50450 | Verify transposon insertion |
| Pa229[#] | gctgttcgacggcagtatc | - | Up PA14_19120 | Sanger sequencing |
| Pa230 | ggggacaagtttgtacaaaaaagcaggctcaaactgcaacgctttctcgat | *attB1* | Up PA14_19120 | Generate mutant |
| Pa231 | tacgcttcagatgaggcccagcaaaaagcctccgtcattcct | - | Up and down PA14_19120 | Generate mutant |
| Pa232[#] | aacggctgacgacctcac | - | Down PA14_19120 | Sanger sequencing |
| Pa233 | aggaatgacggaggctttttgctgggcctcatctgaagcgta | - | Up and down PA14_19120 | Generate mutant |
| Pa234 | ggggaccactttgtacaagaaagctgggtagtggtagcgcaacagcatct | *attB2* | Down PA14_19120 | Generate mutant |
| Pa312 | ggggacaagtttgtacaaaaaagcaggctcattctacctgctcaacagccg | *attB1* | Up PA14_45960 | Generate mutant |
| Pa313 | agaggcaagatcagagagtaataagaccgtcaaccaaggccatagc | - | Up and down PA14_45960 | Generate mutant |
| Pa314 | gctatggccttggttgacggtcttattactctctgatcttgcctct | - | Up and down PA14_45960 | Generate mutant |
| Pa315 | ggggaccactttgtacaagaaagctgggtagaaacggctgagttcccaga | *attB2* | Down PA14_45960 | Generate mutant |
| Pa316[#] | tcaacatggtcacctccagc | - | Up PA14_45960 | Sanger sequencing |
| Pa317[#] | tccagcgtacagtcggaaaag | - | Down PA14_45960 | Sanger sequencing |

| | | | | |
|---|---|---|---|---|
| Pa339 | gggg<u>acaagtttgtacaaaaaagcaggctca</u>ggacgaagagaccctgctga | *attB1* | Up PA14_05360 | Generate mutant |
| Pa340 | cctatgctcaggcctgctctcttcatatttggcccccgc | - | Up and down PA14_05360 | Generate mutant |
| Pa341 | gcgggggccaaatatgaagagagcaggcctgagcatagg | - | Up and down PA14_05360 | Generate mutant |
| Pa342 | gggg<u>accactttgtacaagaaagctgggta</u>caggtccagcacattcagcc | *attB2* | Down PA14_05360 | Generate mutant |
| Pa343[#] | ccaggtactcaaggccgaga | - | Up PA14_05360 | Sanger sequencing |
| Pa344[#] | tgcctgagtaccccttacgg | - | Down PA14_05360 | Sanger sequencing |
| Pa345 | gggg<u>acaagtttgtacaaaaaagcaggctca</u>gggtgagggtctcgaggatc | *attB1* | Up PA14_05320 | Generate mutant |
| Pa346 | gcggccggatatcaggaaacctgttccatgttcgccctata | - | Up and down PA14_05320 | Generate mutant |
| Pa347 | tatagggcgaacatggaacaggtttcctgatatccggccgc | - | Up and down PA14_05320 | Generate mutant |
| Pa348 | gggg<u>accactttgtacaagaaagctgggta</u>tgcatgccgaacacctcatc | *attB2* | Down PA14_05320 | Generate mutant |
| Pa349[#] | cgccgtcgatcatcaggatg | - | Up PA14_05320 | Sanger sequencing |
| Pa350[#] | tcgaggaagccctggtgttg | - | Down PA14_05320 | Sanger sequencing |
| Pa351 | gggg<u>acaagtttgtacaaaaaagcaggctca</u>cgataaccgtggcaaggtcc | *attB1* | Up PA14_16500 | Generate mutant |
| Pa352 | gcggcaccggctgttcgtgcatgtttctctccggga | - | Up and down PA14_16500 | Generate mutant |
| Pa353 | tcccggagagaaacatgcacgaacagccggtgccgc | - | Up and down PA14_16500 | Generate mutant |
| Pa354 | gggg<u>accactttgtacaagaaagctgggta</u>cgtactcgcccttgatgtgc | *attB2* | Down PA14_16500 | Generate mutant |
| Pa355[#] | tcatggcggaaagtccttgc | - | Up PA14_16500 | Sanger sequencing |
| Pa356[#] | tcatcgcggtgtccttgttg | - | Down PA14_16500 | Sanger sequencing |
| Pa357 | gggg<u>acaagtttgtacaaaaaagcaggctca</u>ggagagctggacctgatcgt | *attB1* | Up PA14_55160 | Generate mutant |
| Pa358 | atcggcgttcagcaggctgtcgcagtcataagtgatggc | - | Up and down PA14_55160 | Generate mutant |
| Pa359 | gccatcacttatgactgcgacagcctgctgaacgccgat | - | Up and down PA14_55160 | Generate mutant |
| Pa360 | gggg<u>accactttgtacaagaaagctgggta</u>cgggcccgagaaaaatcctc | *attB2* | Down PA14_55160 | Generate mutant |
| Pa361[#] | gaaccgcctgctggtcac | - | Up PA14_55160 | Sanger sequencing |
| Pa362[#] | tgtgtcgttcttgtcactcgt | - | Down PA14_55160 | Sanger sequencing |

| | | | | |
|---|---|---|---|---|
| Pa367 | tcgaccagcaggacatcaac | - | Up PA14_50480 | Verify transposon insertion |
| Pa368 | atgctcatgctctactccgc | - | Down PA14_50480 | Verify transposon insertion |
| Pa369 | agggtagagtcagccggaat | - | Up PA14_58750 | Verify transposon insertion |
| Pa370 | gccttcaccagcatagggtt | - | Down PA14_58750 | Verify transposon insertion |
| Pa371 | caatcggtgatctcgcagacc | - | Up PA14_05190 | Verify transposon insertion |
| Pa372 | aggaaatccagttggccgag | - | Down PA14_05190 | Verify transposon insertion |
| Pa373 | actggatggggccgatgaa | - | Up PA14_05180 | Verify transposon insertion |
| Pa374 | ataccatgcccaccaggttg | - | Down PA14_05180 | Verify transposon insertion |
| Pa381 | ggggacaagtttgtacaaaaaagcaggctcacgaagtgatctacccgacgg | *attB1* | Up PA14_64230 | Generate mutant |
| Pa382 | gtcgctgccctcaggagatccgaagccgtaccac | - | Up and down PA14_64230 | Generate mutant |
| Pa383 | gtggtacggcttcggatctcctgagggcagcgac | - | Up and down PA14_64230 | Generate mutant |
| Pa384 | ggggaccactttgtacaagaaagctgggtagatgaagatgtagcggccga | *attB2* | Down PA14_64230 | Generate mutant |
| Pa385[#] | aggaggccagcttcatcgt | - | Up PA14_64230 | Sanger sequencing |
| Pa386[#] | gcagacgaacagacccatca | - | Down PA14_64230 | Sanger sequencing |
| Pa387 | ggggacaagtttgtacaaaaaagcaggctcactgatcctgttccagcagca | *attB1* | Up PA14_30650 | Generate mutant |
| Pa388 | atctagctggcggcatcgac(aatcacgctgcacctcgtc) | - | Up and down PA14_30650 | Generate mutant |
| Pa389 | gacgaggtgcagcgtgatt(gtcgatgccgccagctagat) | - | Up and down PA14_30650 | Generate mutant |
| Pa390 | ggggaccactttgtacaagaaagctgggtattgcagcgcttgatctggta | *attB2* | Down PA14_30650 | Generate mutant |
| Pa391[#] | ctgatcctgttccagcagca | - | Up PA14_30650 | Sanger sequencing |
| Pa392[#] | cctgctccatgccgacg | - | Down PA14_30650 | Sanger sequencing |
| Pa393 | ggggacaagtttgtacaaaaaagcaggctcacaagtacaacgttccgctgc | *attB1* | Up PA14_52570 | Generate mutant |

| | | | | |
|---|---|---|---|---|
| Pa394 | ttaatggtttggctcttgatctttcattcctttctcctcacgcgaat | - | Up and down PA14_52570 | Generate mutant |
| Pa395 | attcgcgtgaggagaaaggaatgaaagatcaagagccaaaccattaa | - | Up and down PA14_52570 | Generate mutant |
| Pa396 | ggggaccactttgtacaagaaagctgggtaaaactgctctaccgccttcc | *attB2* | Down PA14_52570 | Generate mutant |
| Pa397[#] | ggcgtctacaccaccgatc | - | Up PA14_52570 | Sanger sequencing |
| Pa398[#] | aaactgctctaccgccttcc | - | Down PA14_52570 | Sanger sequencing |
| Pa399 | ggggacaagtttgtacaaaaaagcaggctcactcgatgatgatgccgagct | *attB1* | Up PA14_58730 | Generate mutant |
| Pa400 | tcttttcagcattagcctattagcgctgagctttcatgtatctctccattg | - | Up and down PA14_58730 | Generate mutant |
| Pa401 | caatggagagatacatgaaagctcagcgctaataggctaatgctgaaaag a | - | Up and down PA14_58730 | Generate mutant |
| Pa402 | ggggaccactttgtacaagaaagctgggtaccaattgggtctgtagcggt | *attB2* | Down PA14_58730 | Generate mutant |
| Pa403[#] | cgttcggagatatccaggcc | - | Up PA14_58730 | Sanger sequencing |
| Pa404[#] | ctacatctccatcggcaccc | - | Down PA14_58730 | Sanger sequencing |
| Pa405 | ggggacaagtttgtacaaaaaagcaggctcatccctatgtgcgcgaatacc | *attB1* | Up PA14_50290 | Generate mutant |
| Pa406 | cagctggttggcctgggcaagggccatggtgatttcc | - | Up and down PA14_50290 | Generate mutant |
| Pa407 | ggaaatcaccatggcccttgcccaggccaaccagctg | - | Up and down PA14_50290 | Generate mutant |
| Pa408 | ggggaccactttgtacaagaaagctgggtaatcggtgtaatcggtcgagc | *attB2* | Down PA14_50290 | Generate mutant |
| Pa409[#] | tccctatgtgcgcgaatacc | - | Up PA14_50290 | Sanger sequencing |
| Pa410[#] | cgacggagatgttcagcgta | - | Down PA14_50290 | Sanger sequencing |
| Pa464 | ggggacaagtttgtacaaaaaagcaggctcagagtcagccggaatattacc ca | *attB1* | Up PA14_58750 | Generate mutant |
| Pa465 | attagtccttggtcacgcggtcgttcattgggagtggtcg | - | Up and down PA14_58750 | Generate mutant |
| Pa466 | cgaccactcccaatgaacgaccgcgtgaccaaggactaat | - | Up and down PA14_58750 | Generate mutant |
| Pa467 | ggggaccactttgtacaagaaagctgggtaggcaactcggcaccaaaatt | *attB2* | Down PA14_58750 | Generate mutant |
| Pa468[#] | cagtcaatagagccagtcacac | - | Up PA14_58750 | Sanger sequencing |
| Pa469[#] | ggtagtggatagcgttcggg | - | Down PA14_58750 | Sanger sequencing |
| Sa046 | ctcgattctattaacaaggg | - | "Upstream" *bursa aurealis* transposon | Verify transposon insertion |

| | | | | |
|---|---|---|---|---|
| Sa047 | gcttttctaaatgttttttaagtaaatca | - | “Buster” *bursa aurealis* transposon | Verify transposon insertion |
| Sa094 | agaaaagcctatggaaattgccctc | - | Up SAUSA300_1992 | Verify transposon insertion |
| Sa095 | tcaccgatgcatagcagtgt | - | Down SAUSA300_1992 | Verify transposon insertion |
| Sa096 | gtataatgacagtgaggagagtggtg | - | Up SAUSA300_1989 | Verify transposon insertion |
| Sa097 | aggacgcgctatcaaacatt | - | Down SAUSA300_1989 | Verify transposon insertion |
| Sa098 | gagagtgtgatagtaggtggaattat | - | Up SAUSA300_1991 | Verify transposon insertion |
| Sa099 | gcgtggtatatcatcagcgc | - | Down SAUSA300_1991 | Verify transposon insertion |

* Up = upstream arm of gene; down = downstream arm of gene.

[#] Primers utilized for Sanger sequencing.

^ Site is underlined in primer sequence.
