## Extended Figures for "Interspecies surfactants serve as public goods enabling surface motility in *Pseudomonas aeruginosa*"

**TITLE**

Running head: Surfactants enable motility in *P. aeruginosa*

*Delayna L. Warrell and Tiffany M. Zarrella contributed equally to this work. Author order was determined alphabetically.

Key words: *Pseudomonas aeruginosa*, *Staphylococcus aureus*, surfactants, motility, polymicrobial interactions, phenol-soluble modulins

The authors declare no conflict of interest.

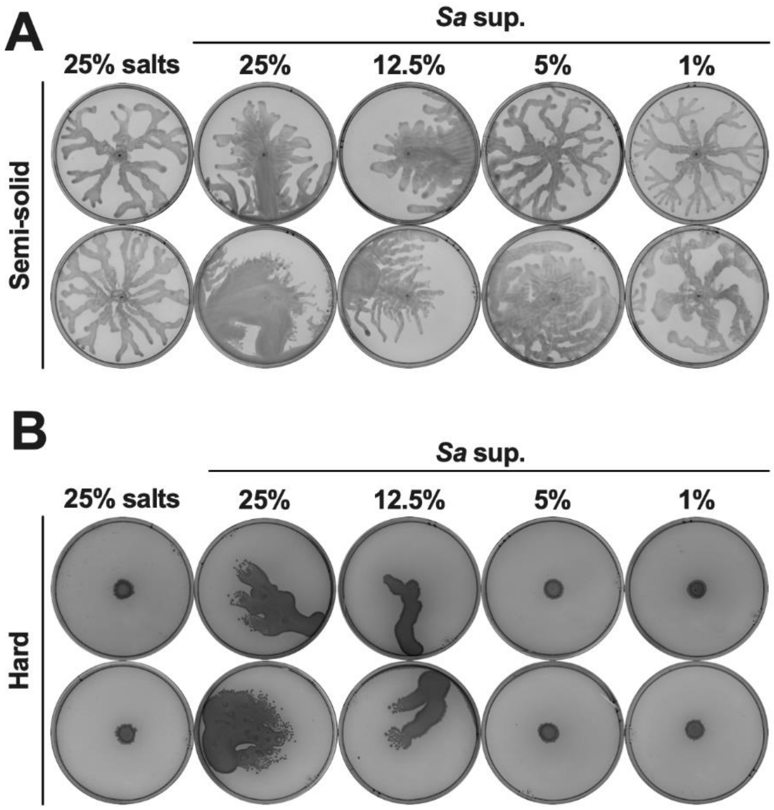


**Extended Figure 1. *S. aureus* secreted products enable *P. aeruginosa* surface spreading in a dose-dependent manner.** Additional replicates shown for Figure 1.

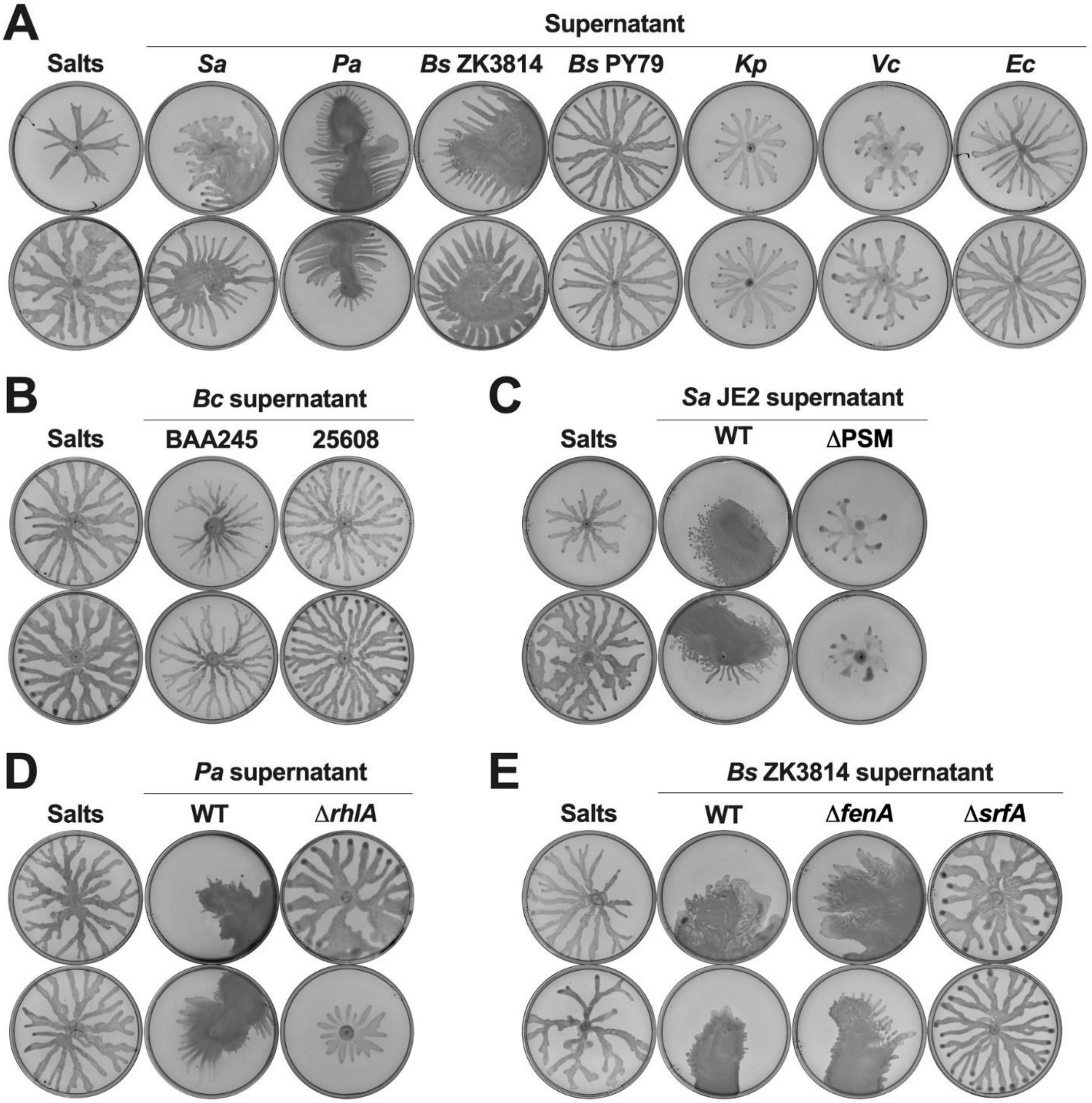


**Extended Figure 2. Interspecies secreted surfactants facilitate surface spreading in *P. aeruginosa*.** Additional replicates shown for Figure 2.

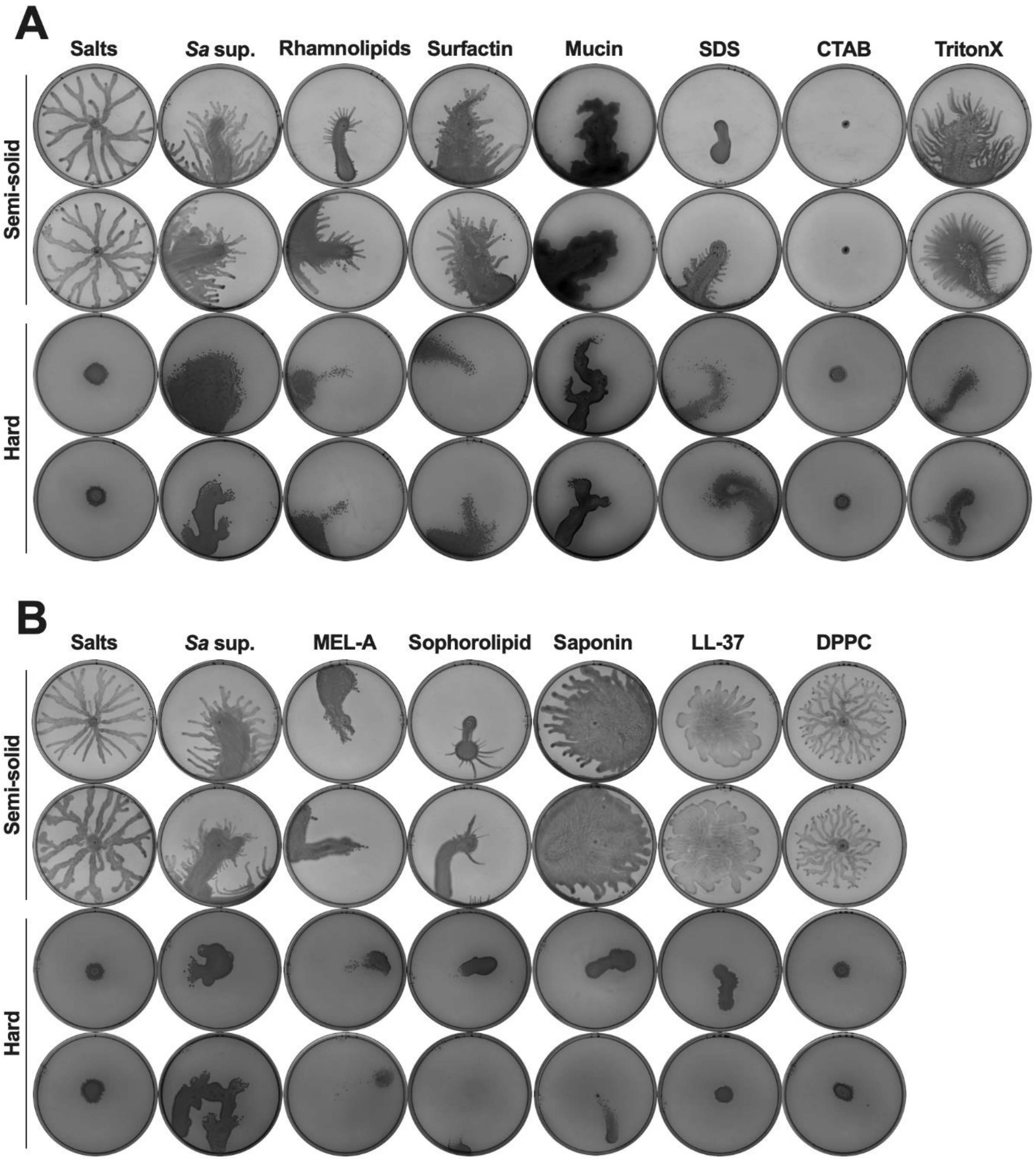


**Extended Figure 3. The addition of diverse biotic and synthetic surfactants is sufficient to enable surface spreading in *P. aeruginosa*.** Additional replicates shown for Figure 3.

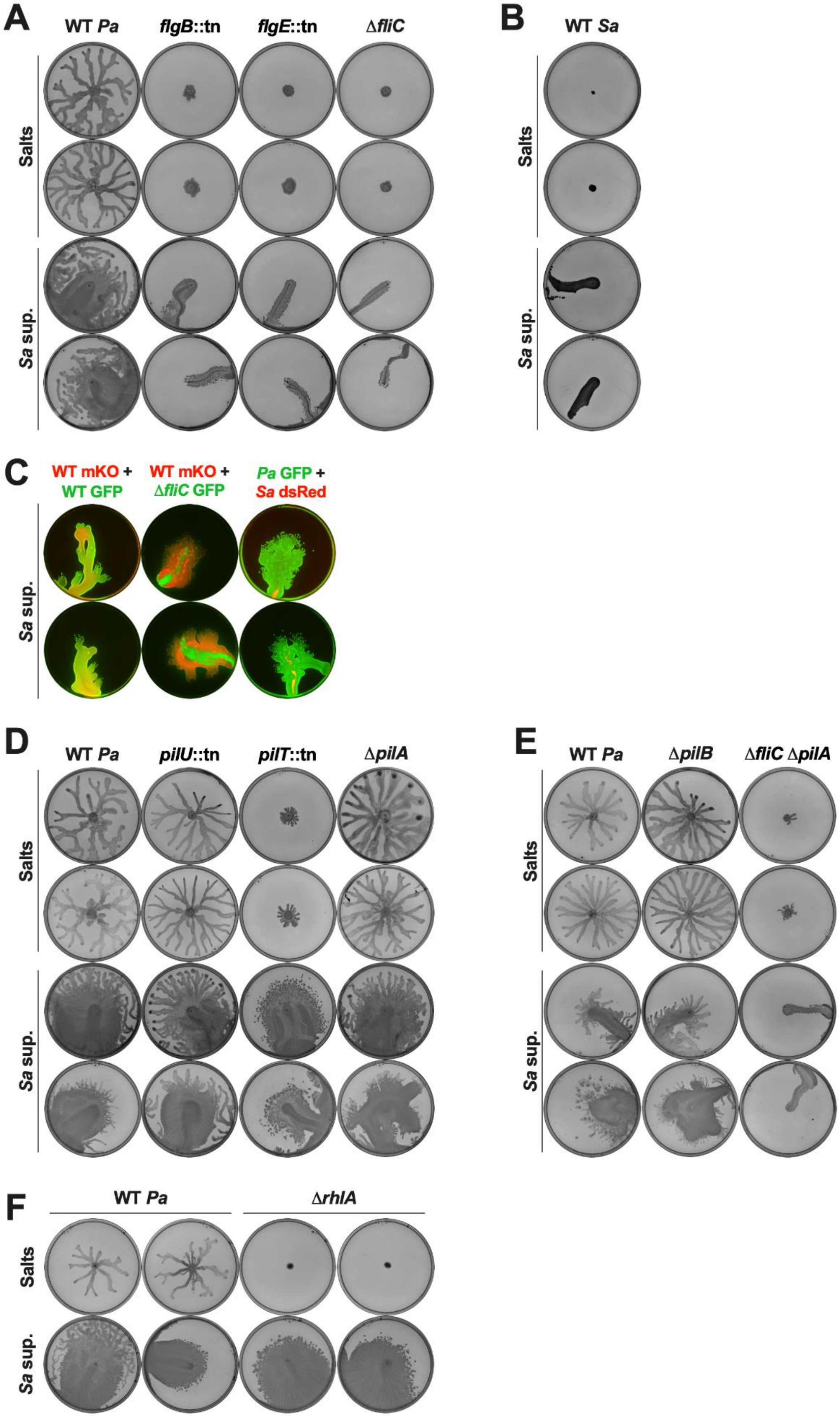
A
WT Pa
flgB::tn
flgE::tn
ΔfliC
Salts
Sa sup.
B
WT Sa
Salts
Sa sup.
C
WT mKO + WT GFP
WT mKO + ΔfliC GFP
Pa GFP + Sa dsRed
Sa sup.
D
WT Pa
pilU::tn
pilT::tn
ΔpilA
Salts
Sa sup.
E
WT Pa
ΔpilB
ΔfliC ΔpilA
Salts
Sa sup.
F
WT Pa
ΔrhlA
Salts
Sa sup.

**Extended Figure 4. Surface spreading requires flagella, but not pili or rhamnolipids.**

Additional replicates shown for Figure 4.

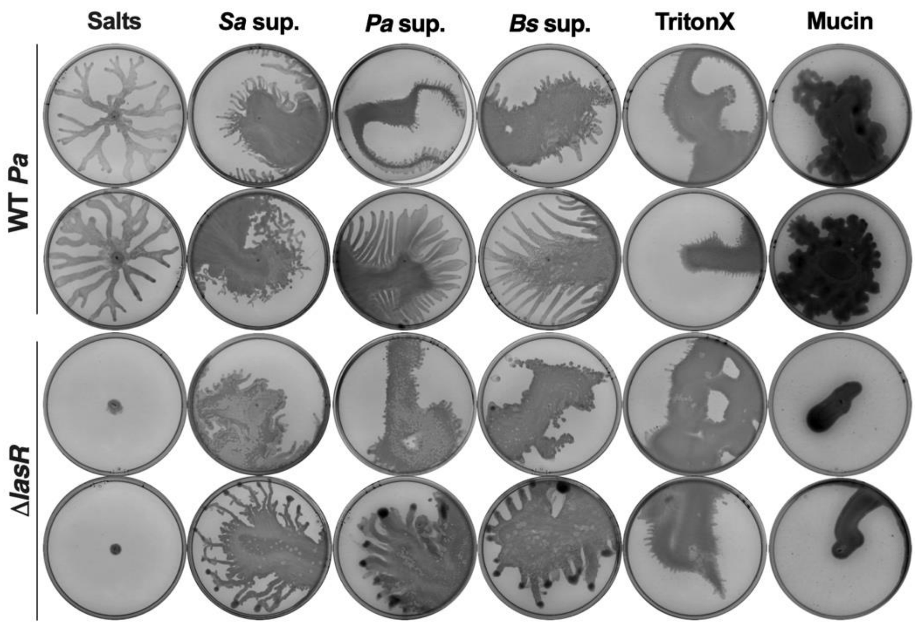


**Extended Figure 6. The Las system is required for motility on mucin, but not on surfactants.** Additional replicates shown for Figure 6.
